# Spatial Deconvolution of HER2-positive Breast Tumors Reveals Novel Intercellular Relationships

**DOI:** 10.1101/2020.07.14.200600

**Authors:** Alma Andersson, Ludvig Larsson, Linnea Stenbeck, Fredrik Salmén, Anna Ehinger, Sunny Wu, Ghamdan Al-Eryani, Daniel Roden, Alex Swarbrick, Åke Borg, Jonas Frisén, Camilla Engblom, Joakim Lundeberg

## Abstract

In the past decades, transcriptomic studies have revolutionized cancer treatment and diagnosis. However, tumor sequencing strategies typically result in loss of spatial information, critical to understand cell interactions and their functional relevance. To address this, we investigate spatial gene expression in HER2-positive breast tumors using Spatial Transcriptomics technology. We show that expression-based clustering enables data-driven tumor annotation and assessment of intra-and interpatient heterogeneity; from which we discover shared gene signatures for immune and tumor processes. We integrate and spatially map tumor-associated types from single cell data to find: segregated epithelial cells, interactions between B and T-cells and myeloid cells, co-localization of macrophage and T-cell subsets. A model is constructed to infer presence of tertiary lymphoid structures, applicable across tissue types and technical platforms. Taken together, we combine different data modalities to define novel interactions between tumor-infiltrating cells in breast cancer and provide tools generalizing across tissues and diseases.

## INTRODUCTION

Breast cancer is a vile disease, every day claiming more than a hundred lives in the US alone and inducing tremendous suffering among those affected.[1] To better understand, diagnose, and treat breast cancer, extensive studies into the genomic underpinnings of the disease have been conducted; one outcome of this work is the establishment of several clinically relevant subtypes. These breast cancer subtypes exhibit varying characteristics, including drug sensitivities, and as a consequence, partially dictate the choice of treatment strategy.[2–4] One of the major breast cancer subtypes is defined by an enrichment of the HER2 (human epidermal growth factor receptor 2) expression on tumor cells, often caused by the amplification of a region on chromosome 17 (cytogenetic band chr17q12) comprising the HER2 (*ERBB2*) gene; these tumors are referred to as *HER2-positive* tumors.[5,6] An estimated 15-20% of all breast cancers tumors are HER2-positive, and these tumors often exhibit aggressive growth and demand intense treatment.[7,8] Understanding the molecular processes responsible for tumor growth and development has shown to be of utter importance; exemplified by the drastic improvement in patients’ prognosis after the introduction of HER2-targeted therapies.[9] Despite this, many patients with HER2-positive breast tumors still succumb to the disease. Therefore, to deliver more effective treatments and develop tools for early detection, we must continue to chart the disease’s molecular profile from new and unexplored perspectives.

Tumors share an intimate relationship with their surroundings, and are best studied accordingly; i.e., in the context of their environment. For example, the success of immunotherapy in tumor treatment stems from its ability to interfere with interactions between cancer and certain immune cells.[10] In addition, the presence of sites promoting cell-cell interactions, such as tertiary lymphoid structures (TLSs), have shown to hold predictive power of treatment outcome in HER2-positive tumors.[11–13] Evidently, the characterization of the cell type population within tumors and their environment is imperative to understand the disease.

Techniques such as immunohistochemistry (IHC), *in situ* hybridization (ISH) and single cell RNA sequencing (scRNA-seq) have been rigorously used to study the transcriptome as well as the physical location of genes and proteins.[14–16] However, these methods are usually burdened by a tradeoff between spatial resolution and throughput. To exemplify, techniques such as IHC and ISH may provide high spatial resolution, but tend to require *a priori* selection of targets, making these methods less suited for exploratory analysis. In contrast, scRNA-seq offers insights into the transcriptome of cells in the higher order of magnitudes, but their spatial origin is to a large extent lost.[17] Spatial Transcriptomics (ST), as described by Ståhl and Sálmen et.al., presents a solution to this dilemma, by providing spatially resolved and transcriptome-wide expression information.[18] Cell interactions and spatial context are key components of the tumor ecosystem, however this space is inhabited by a diverse population of complex cell types that cannot be defined by a few marker genes or surface receptors; hence the benefits of using a technique like ST.[19,20] Although ST does not provide single-cell resolution, this issue can be addressed by leveraging information from scRNA-seq, spatially mapping cell types or clusters by integration of the two data modalities.[21,22]

In this study, we used ST to survey the spatial patterns of gene expression and cell types in 36 samples collected from eight HER2-positive individuals. Intra-and inter-patient heterogeneity was examined using a number of different methods, including expression-based clustering and single cell data integration. We here present novel findings of expression signatures shared among HER2-positive patients, cell type co-localization patterns and a predictive model for TLS-sites.

## RESULTS

HER2-positive tumors from eight individuals (patient A-H) were subjected to ST with three alternatively six sections collected from each tumor, this resulted in a total of 36 ST-sections (See Supplementary Figure 1 for details regarding the experimental setup). For brevity, we will refer to sections originating from the same individual as replicates. Figure 1 provides an overview of the analysis workflow and methods used for this purpose.

**Figure 1.**
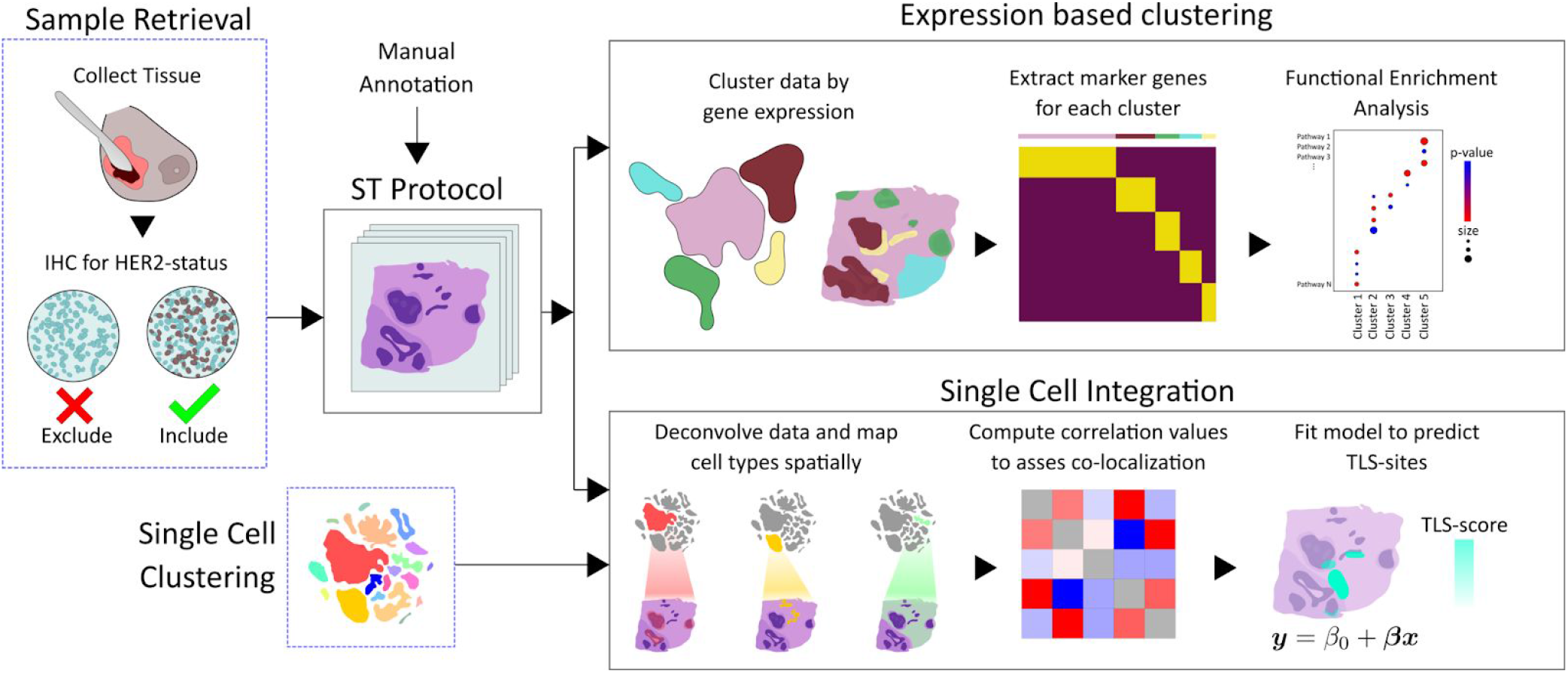
Overview of study. After sample retrieval, we performed ST on 36 sections confirmed as being HER2-positive. A trained pathologist manually annotated one section from each sample. Expression-based clustering and single cell data integration was applied to explore the spatial expression profiles and cell type interactions in our data. Marker genes were extracted for each of the clusters and subjected to functional enrichment analysis, which allowed us to biologically annotate them. By deconvolving the cell types in each spot we could infer novel patterns of cell type co-localization and design a model for prediction of TLSs. Blue dashed boxes indicate steps in the process executed by external groups or individuals.

### Manual annotation

One section from each tumor was examined and annotated by a pathologist. Based on morphology, regions were labeled as either: *in situ* cancer, invasive cancer, adipose tissue, immune infiltrate, or connective tissue, see Figure 2A and Supplementary Figure 3; not all regions were represented in every patient.

**Figure 2.**
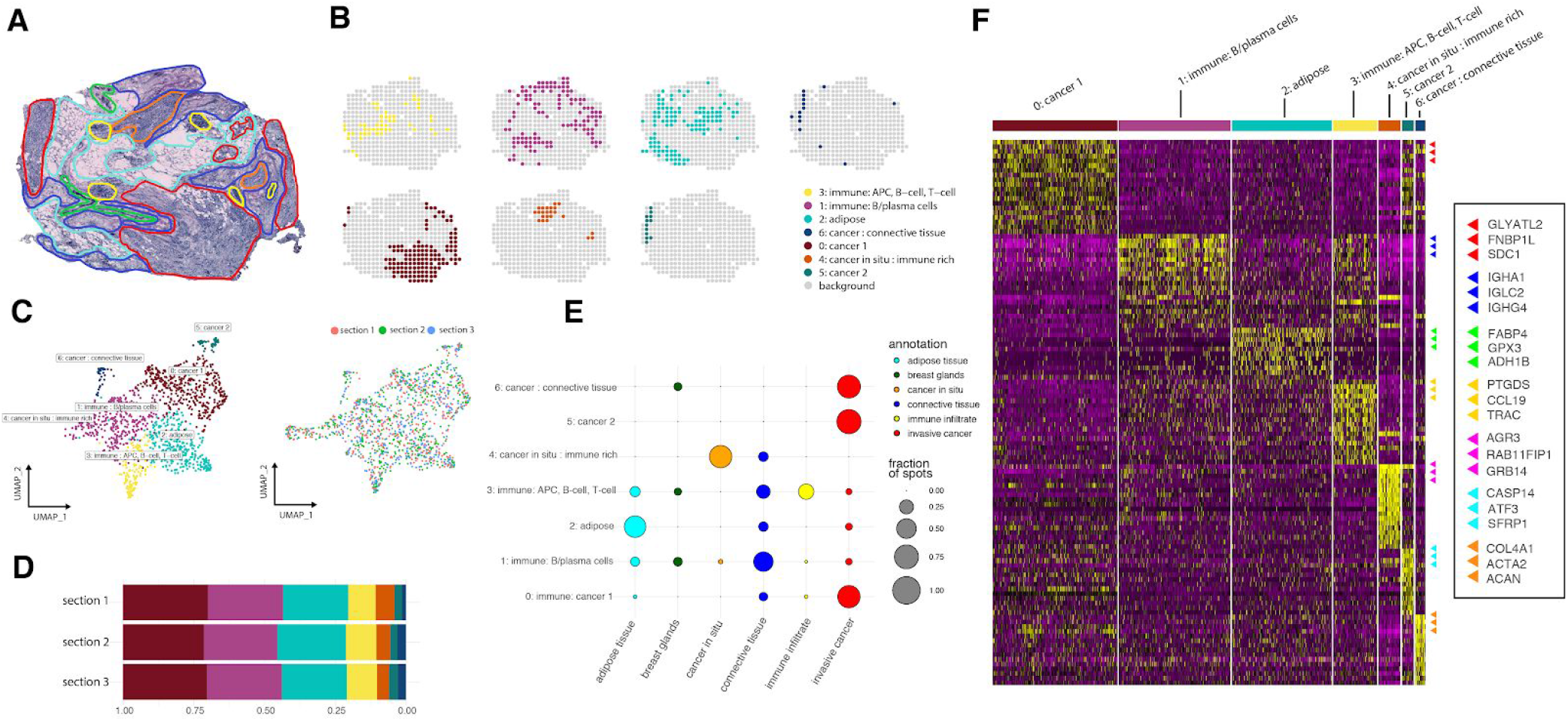
Clustering of spatial data. **A.** Morphological regions were annotated by a pathologist into six distinct categories: invasive cancer (red), adipose tissue (cyan), connective tissue (blue), breast glands (green), *in situ* cancer (orange) and inflammatory cells (yellow). **B.** Split view of each cluster’s distribution across one tissue section. **C.** UMAP projection of 446 spots from patient G colored based on cluster identity. **D.** Proportions of spots assigned to each cluster across the three consecutive tissue sections in patient G. **E.** Dot plot showing the overlap between clusters and annotated regions. The size of the dots represent the proportion of cluster spots belonging to an annotated region. The pathologist’s annotations are given on the x-axis, cluster annotations are found on the y-axis. **F.** Heat map of the clusters and the most highly differentially expressed genes for patient G. Each cluster was annotated based on morphological region together with marker genes

### Initial data characterization

Before conducting any downstream analysis, we assessed the data’s character with respect to patients and sections. First, the data was filtered and normalized (see Methods) to prepare it for subsequent analysis. Next, UMAP was applied to the processed data to obtain a two-dimensional embedding, see Supplementary Figure Supplementary Figure 2.[23] Upon inspection of the UMAP-projection, spatial capture locations (hereafter; spots) from the same patient were seen to cluster together, while a high degree of intermixing between spots from different replicates (within each patient) occurred. Interpatient heterogeneity can be expected when working with tumor samples.[24] Thus, to properly capture the nuances of each patient’s molecular profile and not risk quenching weak signals, we analyzed each patient separately, then compared the outcomes.

### Expression-based clustering

In the patient-wise analysis, we split the data into mutually exclusive patient subsets. A normalization procedure, designed to remove technical noise and batch effects, was then applied to the raw data of each patient subset, i.e., all replicates from the same patient were jointly processed. Results from our analyzes, exemplified through patient G, are presented in Figure 2. Equivalent results for all patients are found in Supplementary Figure 3. Next, the normalized expression data was clustered using a shared nearest neighbor (SNN) approach, resulting in groups (clusters) of spots with similar gene expression profiles, see Figure 2C. Spots neighboring in physical space frequently resided near each other in expression space, i.e., they were often assigned to the same cluster, see Supplementary Figure 4. The clusters were to a large extent spatially coherent but, just as tumors or immune infiltrates, not always confined to a single region, see Figure 2B. Cluster arrangement in the physical domain was consistent across replicates (Supplementary Figure 4 and Supplementary Figure 6).

### Cluster Annotation

To better understand what biological entities the expression-based clusters represented, we first contrasted each cluster’s expression profile towards the rest (differential gene expression analysis), resulting in a set of *marker genes* characteristic of each cluster, see Figure 2F. The marker genes were selected by a combined cutoff with respect to their adjusted p-value and fold change, see Methods and Supplementary Data 3. To investigate which biological pathways that the clusters were enriched for, we queried their marker genes against the GO:BP (GO Biological Processes) database using *g:Profiler*.[25] From the set of marker genes and their associated pathways, high-level functional annotations could be assigned to the expression-based clusters, motivations for the cluster annotations are found in Supplementary Data 2. These annotations are by no means exhaustive, but provide guidance in subsequent analysis.

### Comparison of clusters with manually defined regions

We related the regions defined by the pathologist to the expression-based clusters by computing the fraction of spots within a region that belonged to each cluster, shown in Figure 2E. Strong concordance was observed; clusters enriched for immune-related processes overlapped well with the immune-infiltrates, those with cancer associated pathways fell into the cancerous regions, and so forth, see Supplementary Data 10. This comparison established the existence of a relationship between the pathologist’s observations and clusters formed by data driven analysis. Notably, an *in situ* cancer region that consisted of as few as 3 spots correctly clustered with physically separated, but identically annotated, spots (cluster 4 in Figure 2B and Supplementary Figure 5), attesting to the sensitivity of ST and verifying that clusters are not based on physical proximity alone. In patient H, parts of a region labeled as *in situ* cancer were consistently, across all replicates, inhabited by two expression-based clusters; one (cluster 1, patient H) that overlapped with the other cancer areas while the second (cluster 4, patient H) was enriched for immune processes and aligned spatially with the annotated immune infiltrates, see Supplementary Figure 5. These observations suggest that data-driven expression-based clustering captures signals that may be overlooked by visual inspection; thus in some cases providing a more in-depth and nuanced depiction of the tumor tissue. The value of this is manifold; the labour intense and time-consuming process of manual annotation is neither suitable for high throughput analysis, nor is access to a trained pathologist always guaranteed. Importantly, regions where distinct molecular pathways are active can easily be distinguished, something that is hard to do only relying on (human) visual inspection.

### Exploring Intra-and interpatient heterogeneity

Multiple tumor profiles may exist among different individuals, but also within a given patient. To outline this heterogeneity may aid in the design of more nuanced and personalized treatment regimens.[26] Therefore, with the expression-based clusters as a structured framework to operate within, we aimed to assess both intra-and interpatient heterogeneity in our data.

Interestingly, we were able to observe intra-patient heterogeneity at the transcriptome level in most of our patients; in fact, all patients except two (patient B and H) had more than one cluster labeled as cancer. To exemplify, patient E exhibited varying transcription profiles in two spatially separated tumor foci assigned to different clusters (cluster 4 and 5, patient E), see Supplementary Figure 3. While true that such observations may be explained by the practice of “overclustering”, the two clusters were clearly separated and non-neighboring in UMAP-space, which suggested distinct expression profiles. Both clusters had *ERBB2* (encoding the HER2-receptor) listed as a marker gene and were enriched for pathways associated to cell growth, but one of the cluster displayed a high degree of enrichment for immune response related processes (cluster 3, patient E) while apoptotic and regulatory pathways were enriched in the other (cluster 4, patient E), see Supplementary Data 6. These findings implied that one tumor focus (cluster 3, patient E) likely had a higher degree of infiltrating immune cells than the other.

Another example of intrapatient heterogeneity was patient G, with four separate clusters (cluster 0 and 4-6) that were annotated as cancer (see Figure 2), one of them aligned with the two *in situ* cancer regions. Of course, the presence of multiple cancer clusters does not necessarily reflect different tumor types, but rather that the corresponding spatial regions are not homogenous in their expression; for example, as a consequence of hosting distinct immune cell populations. If additional metadata could be obtained for patients, ST might be useful for relating tumor intrapatient heterogeneity to more quantitative metrics, such as survival or treatment response.

### Immune and tumor core signatures

To assess if any universal features were present in our data, we compared clusters across patients with respect to their marker genes. We reasoned that if two clusters shared a large number of marker genes, they should be considered more similar than if not. To translate this notion of similarity into a quantitative metric, we computed the Jaccard Index for every combination of cluster pairs. Five distinct *supergroups (*a.k.a, “clusters of clusters”) emerged after the clusters were hierarchically grouped based on similarity, see Methods and Supplementary Figure 7. Only those supergroups where at least one gene was shared among all members were considered as robust, meaning two supergroups were discarded. In the remaining three supergroups, marker genes present in a majority of the member clusters (at least 80%) were extracted and considered as *core signatures* of the HER2-positive patients. Two core signatures were immune-related: the first being a set of 47 genes, including *APOE* and *C1Q{A,B*,*C}* expressed (but not exclusively) by macrophages, suggesting that clusters in the corresponding supergroup might contain tumor-associated macrophages;[19] the second was a set of 55 genes with lymphocyte and MHC class I-II associated members (e.g., *TRBC1*, *HLA-{A,B}*, *HLA-D{QB1,RA,RB1}*).[27] The third core signature consisted of 11 genes, with several of them being related to cancer and proliferative growth (e.g., *ERBB2, EPCAM* and *CDH1).* [28,29] This cancer core signature was derived from a supergroup where all clusters were annotated as cancer-associated. For clarity, the aforementioned cancer supergroup consisted exclusively of cancer-associated clusters, but not all such clusters were members of this group. See Supplementary Data 1 for the complete core signatures.

### Inference of cell type organization by integration with single cell data

Unsupervised clustering of expression data provides insight into spatial expression motifs present in the data. Still, these expression profiles are generated by cell mixtures consisting of one or more cell types; meaning that a one-to-one relationship between cluster and cell type is not guaranteed. Given how spatial arrangement and patterns of interaction between different cell types have implications for both disease progression and treatment, we wanted to chart each cell type’s spatial distribution within the tissue. For this purpose, we deconvolved our spatial data with scRNA-seq data from five HER2-positive tumors, annotated in three tiers, see Methods.[30] The three tiers are referred to as the *major*, *minor* and *subset* tier (terminology inherited from original source). There were eight different cell types in the major class: myeloid cells, T-cells, B-cells, epithelial cells, plasma cells, endothelial cells, cancer associated fibroblasts (CAFs) and Perivascular Like cells (PVL cells). The minor and subset tiers represented gradually finer partitioning of the major types; with examples such as Macrophages (for brevity Mø) and CD8+ T-cells in the minor level and subsets of these in the lowest tier. Several methods to integrate spatial and scRNA-seq data have been proposed, and success has been shown upon applying these techniques to spatial cancer data.[21] Most of these methods rely on correlation or elevated expression of a select set of marker genes; however, we decided to use a method that takes advantage of the full expression profiles from both data modalities. In short, the method we used (*stereoscope*) decomposes the expression observed in each spatial location — a mixture of transcripts from multiple cells —into contributions from different cell types defined by the single cell data using a probabilistic model.[22] The advantage of this approach is that similar cell types with overlapping sets of marker genes may still be distinguished, since the inference is based on the complete expression profile of each type, which is especially important in a complex and intermixed environment like tumors. Excerpts from the single cell integration analysis are given in Figure 3. Supplementary Data 8 contains visualization of all of the remaining patients and tiers, all output from *stereoscope* is found in Supplementary Data 7.

**Figure 3.**
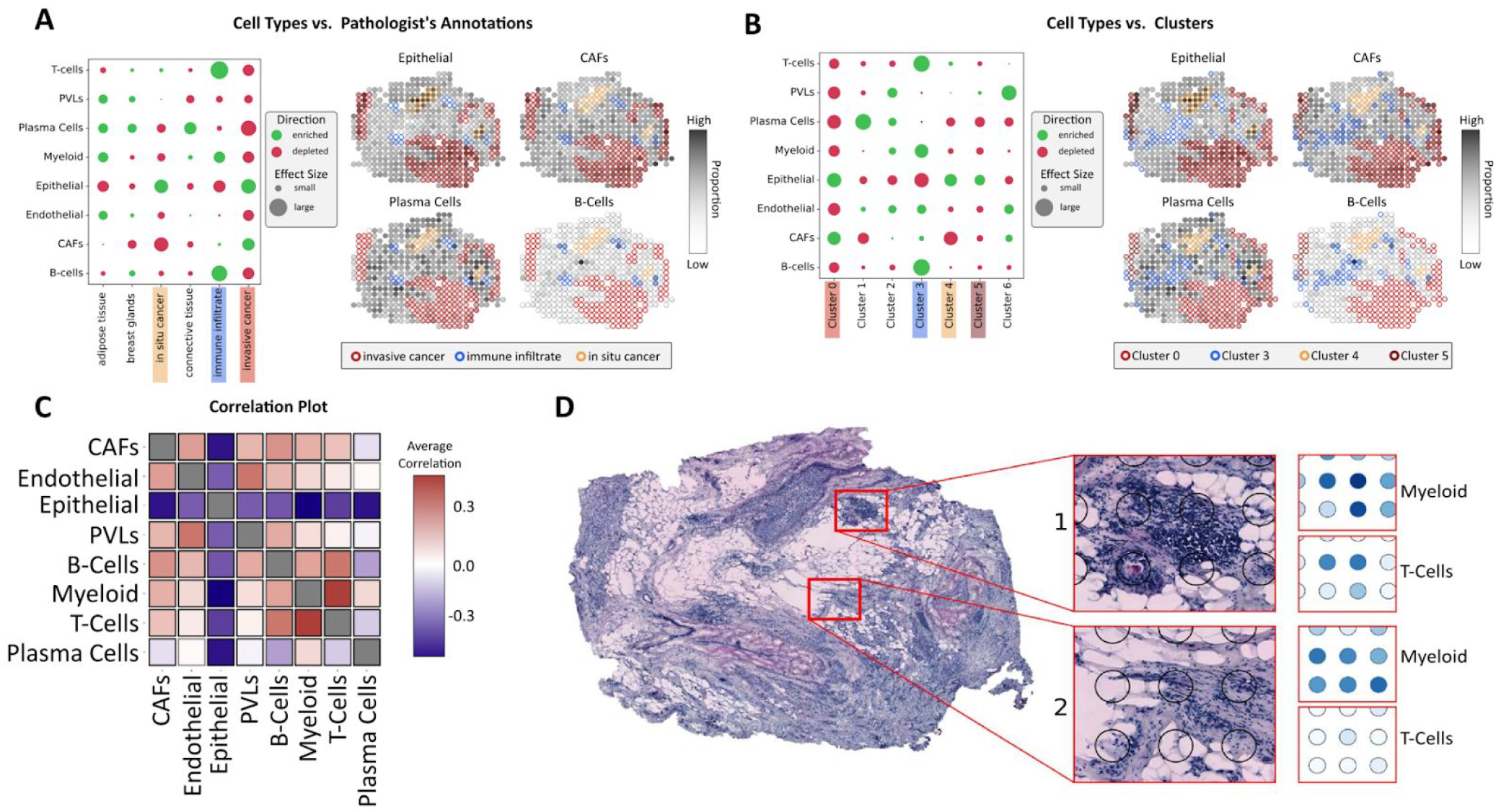
Single cell mapping through *stereoscope*. **A**. Enrichment (green) and depletion (red) of major tier cell types in the regions defined by the pathologist, along with proportion estimates of different cell types (epithelial, CAFs, plasma cells and B-cells). Spots annotated as cancer *in situ*, invasive cancer and immune infiltrate are indicated by border color. **B.** Similar to A but with the regions defined by the expression-based clusters. **C.** Correlation plot of all cell types within the major tier across all sections, a distinct correlation between myeloid cells and T-cells can be observed. **D.** Proportions of myeloid cells and T-cells showing one area with higher (1) respectively lower (2) degree of co-localization. All presented results are associated with patient G, except for subfigure C where correlation values are computed across all patients.

### Enrichment of cell types within manually defined regions

Analogous to the analysis of our expression-based clusters, we examined enrichment/depletion of cell types within the pathology-annotated regions to see how the single cell mapping related to these. We implemented a slightly different approach to assess cell type enrichment, since proportion values represent continuous values in contrast to the discrete cluster labels, see Methods. Several affirmative trends were observed, B and T-cells were enriched in the immune infiltrates while cancer regions showed enrichment of cancer-related cell types and depletion of several cell types, see Figure 3A and Supplementary Data 9. We noted how all patients except patient B showed enrichment of the HER2-related epithelial cancer type in the regions annotated as invasive cancer regions. However, patient B exhibited depletion of the HER2 related type in combination with enrichment of the LUMB type in these areas. Coincidentally, patient B was also unique in having a progesterone-receptor positive profile, in concordance with the LUMB molecular subtype, see Supplementary Figure 14 and Supplementary Table 1.[31] Taken together, these findings were seen as affirmative of our mapping’s validity.

### Enrichment of cell types within expression-based clusters

Curious of which cell types that drove the formation of the expression-based clusters, we conducted the same type of enrichment analysis as for the pathologist’s annotations, but this time we assessed enrichment/depletion of cell types within clusters, see Figure 3B. In patient E — with the two spatially disconnected tumor foci — the cluster associated with apoptotic and regulatory pathways (cluster 4, patient E) was enriched for epithelial cells and depleted of memory B-cells. In contrast the immune rich cancer cluster (cluster 3, patient E) was enriched for memory B cells and CD4+ T-cells, with weaker or no enrichment of cancer types, see Supplementary Figure 11. In Patient G, all clusters annotated as cancer were: depleted of plasma cells, had very low enrichment or were depleted of B and T-cells, and three of four clusters were enriched for epithelial types, see Figure 3. The *in situ* cancer cluster in (cluster 4) was the only cancer cluster in patient G enriched for dendritic cells across all replicates while also being depleted of myofibroblast like CAFs (Cancer Associated Fibroblasts), see Supplementary Figure 13. We also observed how the plasma cell immune cluster (cluster 1, patient G) was enriched for plasma cells while the APC immune cluster (cluster 3, patient G) exhibited stronger enrichment of B-cells, T-cells and myeloid cells. PVL cells were overrepresented in the mixed cancer/connective tissue clusters (cluster 6, patient G), which also showed enrichment of myeloid cells, CAFs and endothelial cells, see Figure 3B and Supplementary Figure 12. Adipocytes or equivalent cell types were not included in the single cell data we used, hence no such types were spatially mapped. Still, the adipose cluster (cluster 2, patient G) showed enrichment of PVL cells, myeloid and plasma cells (albeit weak for the latter two). We expect cell types such as neutrophils and mast cells to be present in our spatial data, given how they infiltrate tumors and are associated with tumor progression; but these types were not present in the single cell data and could therefore not be spatially mapped.[32,33] Agreement with the tissue morphology, pathologist-annotations and expression-based clusters therefore suggests that the single cell data is sufficiently representative of our tissues to provide a reliable mapping of the included types.

### Interactive exploration of results

We have compiled a resource that contains all data and results from the expression-based clustering and single cell mapping; with a graphical user interface (GUI) that enables comprehensive exploration of these, see Code Availability.

### Trends of cell type co-localization

To condense the information generated by the cell mapping and discover putative cellular interactions, we examined the co-localization of types by computing their spot-wise Pearson correlation, see Figure 3C. A positive correlation between two types was considered as indicative of them co-localizing, with the degree of co-localization being proportional to their correlation value; the opposite being true for negative values.

Within the major tier, the most conserved feature, present in all patients, was that epithelial cells anticorrelated with all other types, see Figure 3C. Plasma cells anticorrelated with B-cells in all patients except one (patient A, see Supplementary Figure 8). These findings indicate that B-cells and plasma cells, which differentiate from B-cells, reside at distinct locations within the tumor.[34] It is not clear whether these findings reflect plasma cell migration during local differentiation from tumor-associated B-cells, or if the plasma cells have developed from B-cells outside the tumor microenvironment. Positive correlations of varying strengths between B and T-cells (major tier) were observed in five of the eight patients; patients G and H exhibited particularly strong co-localization signals and their spatial distributions of B and T-cells had an ample overlap, see Figure 5A and Supplementary Figure 9-Supplementary Figure 10. Spatially proximal high densities of B and T-cells may be indicative of TLSs, something we will revisit in later figures and text.[20]

**Figure 5.**
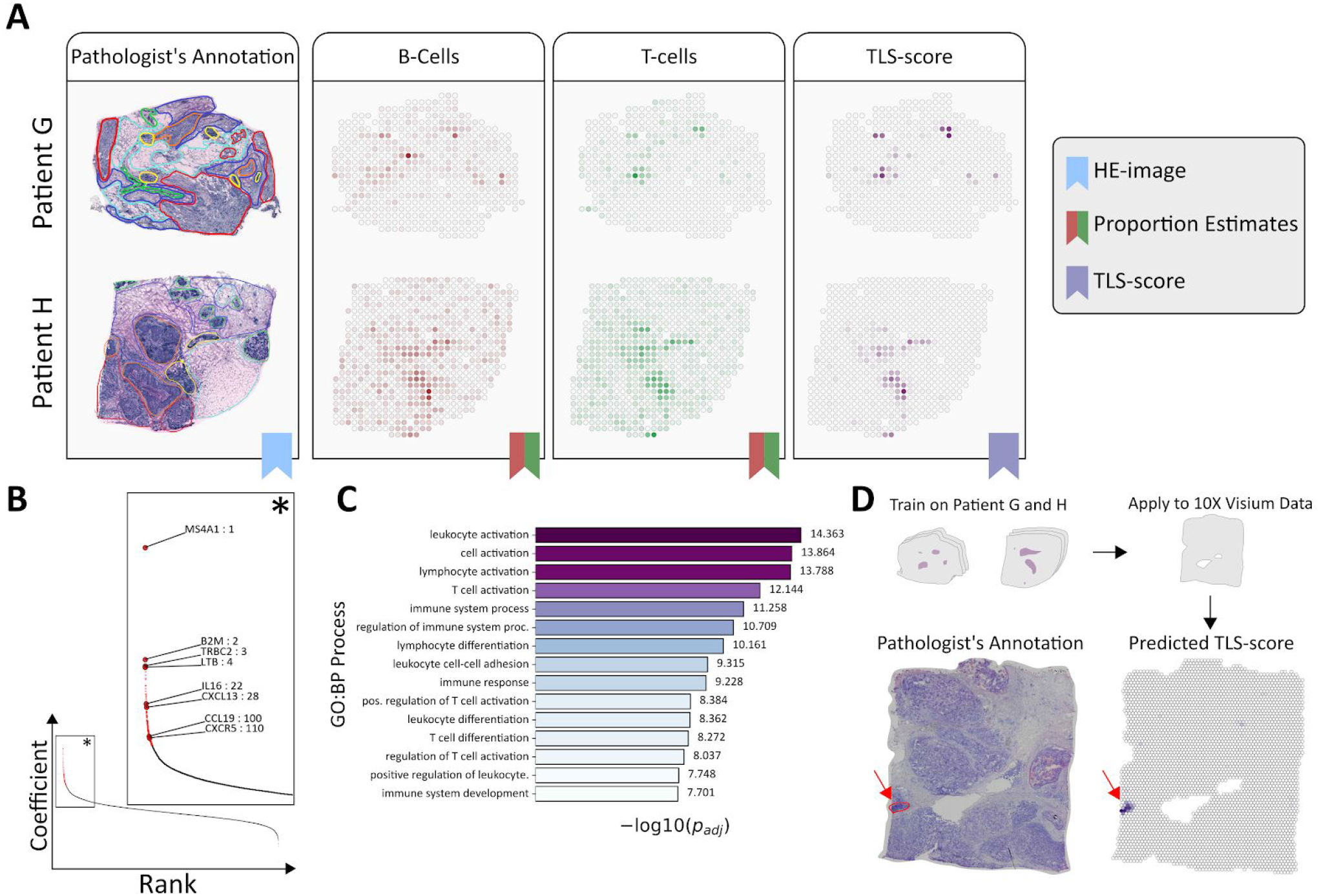
TLS Inference and Prediction. **A.** Proportion estimates of B and T-cells together with the computed TLS-score for patient G and H, annotated HE-images are included for reference. **B.** Rank-plot (coefficient value vs. rank) of the fitted model, genes included in the TLS-signature are indicated by red; excerpt (*) shows the three top ranked genes together with known TLS associated genes. **C.** Top 15 pathways which the TLS-signature showed enrichment of. **D.** Predicted TLS-score for the 10X Visium breast cancer data set, using the model trained on Patient G and H. Pathologist’s annotation for likely TLS-sites (red) are included as a reference.

We also observed that T-cells co-localized with myeloid cells across patients. Interactions between T-cells and myeloid cells are well established and can profoundly affect their respective behavior.[35] Recent studies have also revealed an unprecedented heterogeneity within T-cell and myeloid cell types, where subsets of these exhibited a diverse spectrum of states.[19,36,37] These states present as complex phenotypes not mainly defined by marker genes, but rather the expression profile of a larger gene set. When the finer tiers of these two cell types were examined, several trends of co-localization could be observed; such as weak positive signals between cDC2:CD1C, Mø1:EGR1, and pDC:IRF7 with several CD4+ T-cell populations, including Tfh and Tregs. We also observed a salient correlation between Mø2:CXCL10 macrophages and a T-cell subset (T-cells:IFIT1) across all patients, see Figure 4A, and thus sought an explanation for this.

**Figure 4.**
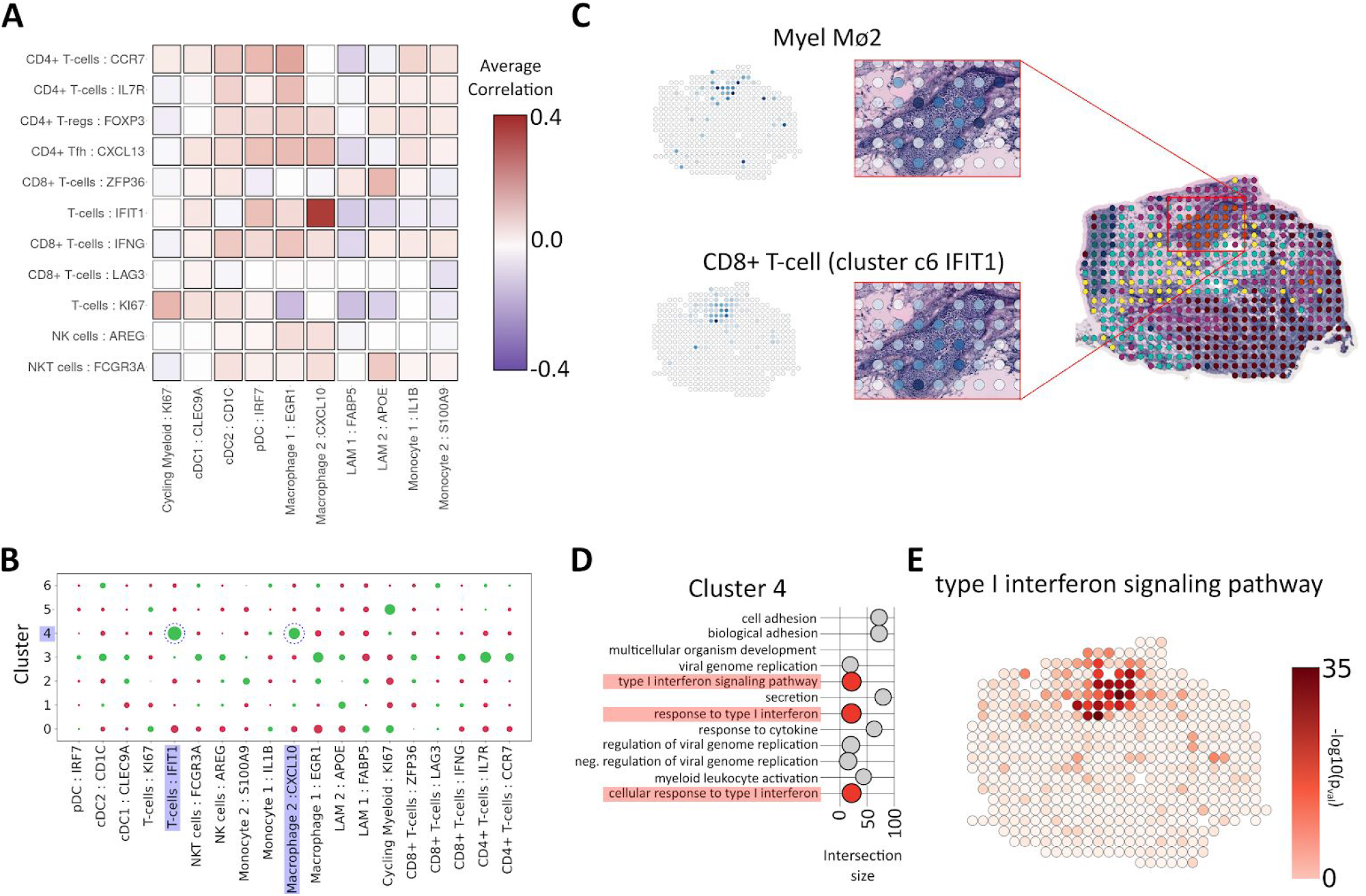
Co-localisation of myeloid cells and T-cells. **A.** Correlation plot of subsets of T-cells and Myeloid cells showing a distinct correlation between the *IFIT1* T-cell subset and Mø2 across all patients. **B.** Enrichment (green) and depletion (red) of subsets of T-cells and Myeloid cells in each manually annotated region, highlighting the presence of the correlated cellypes *IFIT1* T-cell subset and Mø2 within the *in situ* region of patient G. **C.** Proportion estimates for *IFIT1* T-cell subset and Macrophage 2 in patient G, highlighting the annotated *in situ* region. **D.** Pathways enriched by marker genes for cluster 4, interferon signalling pathways are highlighted in red. Intersection size is equivalent to the number of overlapping terms between the marker genes of cluster 4 and the given pathway. **E.** Spot-wise enrichment of type 1 interferon signaling pathway (GO:0060337) visualized on patient G.

### Presence of type I interferon response processes

As indicated in the single cell RNA-seq resource, Mø2:CXCL10 expressed increased levels of the chemoattractants *CXCL10* and *CXCL9.* Both of these chemokines bind *CXCR3*, typically found in T-cells and NK cells.[38,39] Tumor-associated myeloid cells expressing *CXCL9/10* have been described previously and attributed important immunotherapy-induced anti-tumor functions.[19,40–42] Furthermore, the chemokines *CXCL9/10* may be induced by a *type I interferon* stimuli, also reported to be present in the *IFIT1* T-cell subset.[30]

Type I interferon activation within tumors can act directly on tumor cells, to inhibit proliferation or stimulate apoptotic processes, or indirectly by activation of anti-tumor immunity.[43,44] In addition, certain anti-cancer therapies have been shown to induce and depend on type I interferon activation.[44] Given the relevance of type I interferon responses in cancer treatment, we wanted to evaluate whether this process could be associated with Mø2:CXCL10 and IFIT1 T-cell subset co-localization in our spatial data. Thus, we inspected the cell type-within-cluster enrichment results, and noted that a majority of the patients had at least one cluster (e.g., cluster 4 in patient G) enriched for both Mø2:CXCL10 and IFIT1 T-cells, see Figure 4B and Supplementary Data 9. Consequently, we revisited the pathways listed as enriched within the clusters containing both subsets, and noted that type I interferon response related pathways were among the top ranked ones in patient B, E and G, see Figure 4D.

### Spatial enrichment of type I interferon responses

Encouraged by the enrichment of type I interferon signaling pathways in the patient clusters, we conducted a spatial (spot-wise) enrichment analysis, targeted specifically towards these pathways. Briefly summarized, for each spot we determined the intersection between top expressed and pathway-associated genes, to then compute the probability of the overlap occurring by chance; a low probability implies that a spot is enriched for the pathway, see Methods. Regions with high enrichment of the type I interferon response pathways aligned spatially with areas of joint Mø2:CXCL10 and IFIT1 T-cell subset presence, see Figure 4C and 4E. This suggests a spatially restricted type I interferon response in some HER2-positive tumors, which may promote macrophage-induced recruitment of certain T-cells. Further investigations would be useful to establish whether these interactions are relevant to disease outcome.

### Inferring TLSs from cell type proportions

Next, we returned to the patterns of B and T-cell co-localization, and more specifically how this related to TLSs. Our interest in TLSs stems from their cardinal role in antitumor immune responses and relation to clinical outcome as well as treatment response. In the context of cancer, TLSs are one of the locations where tumor antigens are presented to T-cells, promoting a more targeted attack towards the tumors.[20]

Since TLSs are defined by the presence and interaction of multiple cell types, scRNA-seq techniques are suboptimal for studying them unless the sites are separated from the remaining tissue prior to dissociation. We therefore see our use of ST, where each spot represents a small neighborhood populated by multiple cells, as complementary to scRNA-seq methods when studying these structures. While TLSs are not exclusively inhabited by B and T-cells, they are implicated by their joint presence.[45] Having deconvolved the cell type composition of each spot, we were able to identify which spots that exhibited a high degree of co-localization between B and T-cells, ergo potentially constituting parts of a TLS-site. More explicitly, for each spot we computed the probability of two randomly picked cells being a pair of B and T-cells, followed by a subtraction of the average probability of picking any pair within that spot; a metric we dubbed *TLS-score*. A positive TLS-score translates to B and T-cells both being present at a site, negative values the opposite. As expected from the overlap in B and T-cell distribution, patient G and H exhibited small compartmentalized regions with high TLS-score, see Figure 5A, these regions were therefore considered as promising TLS-site candidates.

### Characterizing the gene expression profiles of TLSs

Next, we wanted to assess how the gene expression related to TLS-score. For this purpose we used a simple linear model to predict the TLS-score of a spot based on its (normalized) gene expression, and then extracted the genes with greatest contribution to a positive score (i.e. having large positive coefficient values); we refer to this set of genes as a *TLS-signature*. The number of signature members (171 genes, see Figure 5B and Supplementary Table 2) was determined by a threshold derived from the trained model, see Methods. The three genes with largest coefficient values were: *MS4A1* (a well-known B-cell associated gene, encoding the antigen CD20), *B2M* (encoding a protein that interacts with and stabilizes MHC I) and *TRBC2* (encoding a component of the T-cell receptor). Other signature members have previously been associated with TLSs (e.g., C*XCL13, CXCR5, CCL19* and *LTB)*, see Figure 5B.[20] To see what biological processes the TLS-signature was enriched for, we subjected it to functional enrichment analysis (using g:Profiler, querying against GO:BP). The top processes were all related to cell activation, differentiation and immune response or regulation, see Figure 5C and Supplementary Data 12.

### Predicting presence of TLSs across tissues and platforms

To control for overfitting, we applied the model to (HER2-positive) breast cancer data originating from a different platform (Visium, downloaded from 10xGenomics™ website,[46]). Strong localized signals were observed in the Visium data, overlapping with the region identified as a likely TLS-site by a pathologist, see Figure 5D. Finally, we evaluated the model’s performance on three different types of tissue (rheumatoid arthritis, developmental heart and melanoma), where the results agreed with previous annotations and expectations, see Supplementary Section 1. This suggests that the signature and model are not only representative of our data, but have a more general character. Although further studies are necessary to confirm our findings, charting the molecular profiles of TLSs in this manner could potentially reveal novel therapeutic targets for drugs aiming to promote anticancer immune responses.

## DISCUSSION

Using the Spatial Transcriptomics (ST) technique we have studied eight HER2-positive tumors from a largely unexplored perspective, namely that of their spatial gene expression profiles. Below, we will briefly recapitulate on some of our analyses and their respective ramifications.

We clustered the data based on gene expression and were able to discern sets of genes that distinguished the clusters (i.e., marker genes), which in conjunction with functional enrichment analysis were used to annotate our clusters. From the marker genes, we derived immune and tumor core signatures, elements helpful in attempts to discover new therapeutic targets or alternative treatments.

In addition to expression-based clustering, we used an integrative method to map types found in single cell data onto our samples. This informed us of the types’ spatial distribution within the tissue, and enabled further analysis. When mapped, the cell types arranged as expected; gauged by visual inspection and quantitative measures of enrichment/depletion. From the spatial co-localization patterns we concluded that: the epithelial cancer types tend to be dominant when present (anticorrelating with all other types); plasma cells seemed to spatially segregate from B-cells, in several patients B and T-cells co-localized; a type I interferon associated coupling between certain T-cell and macrophage subsets existed. The proximal location between the *CXCL9/10*-expressing macrophages and T-cells suggests that Mø2 could be recruiting the IFIT1 expressing T-cells into specific locations, which may have implications for the design of future treatment strategies. Still, more extensive efforts are required to properly confirm this synergistic relationship between the two types.

We believe the single cell data to be representative of our tissues, but not perfectly matched, and hence there might be cell types present in the spatial data which we lack the ability to infer proportions of; a limitation to any method relying on external data. For example, neutrophil and mast cells were not included in this analysis, even though they may be present in the samples. Furthermore, some cell types were excluded from the analysis due to low numbers in the single cell data, despite being of biological interest (e.g., one of the dendritic subsets). [19,47–49] There is also a risk of certain cell types being dominant in some regions, meaning that the majority of captured transcripts therewithin originate from this cell type; as a consequence very weak signals from other types may be masked and their presence not properly accounted for. We thus consider the integrative analysis complementary to our expression-based clustering, where the latter is able to discern new motifs of expression while the former allows us to assess the spatial arrangement of known cell types.

The deconvolved spatial data allowed us to identify potential TLS-sites, by studying the joint distribution of B and T-cells. We trained a linear model to identify TLSs based on gene expression, from which we extracted a gene signature associated with potential TLS-sites. As expected for a relevant signature, several of its members had previously been attributed important roles in TLS formation and function. Despite its simplicity, the model generalized across techniques and tissues.

In future studies we envision that cell co-localization patterns may be linked to patient outcome, used to assess drug responses in a spatially restricted manner within tumors, and study functional interactions. For example, there’s an unmet clinical need to understand what dictates how a patient will respond to anti-cancer immunotherapy; for which biomarkers currently used in the clinics are not adequate. To conclude, our study provides new tools and biological insights into the spatial organization in HER2-positive breast cancer tumors, which may help to better understand the underlying disease mechanisms and open up for new vantage points for therapy.

## ACKNOWLEDGEMENTS

We want to thank Patrik Ståhl for valuable comments and advice throughout the process of this study. Furthermore, Jari Häkkinen and Johan Vallon-Christersson provided feedback and comments during the initial phases of this project, which we appreciate tremendously. This work was supported by the Knut and Alice Wallenberg Foundation, Swedish Cancer Society, Swedish Foundation for Strategic Research, the Swedish Research Council, Tobias Stiftelsen, Torsten Söderbergs Foundation, the European Union’s Horizon 2020 research and innovation programme under the Marie Sklodowska-Curie grant agreement No 844712 (CE) and Science for Life Laboratory. We also thank the National Genomics Infrastructure (NGI), Sweden for providing infrastructural support.

## AUTHOR CONTRIBUTIONS

A.A. and C.E. wrote the manuscript with input from the remaining authors. L.S. carried out the laboratory experiments and wrote the corresponding Methods part. A.A., C.E. and L.L performed data analysis, J.H.A.E. (referred to as the pathologist) inspected the patient samples and performed morphological annotation. Å.B. provided samples and assessment of the biological findings. J.L., J.F., F.S. and Å.B. planned the spatial transcriptomics study. S.W., G.A.E., D.R, and A.S. provided early access to the single cell data used in the study as well as guidance regarding how to orient interpret the results related to it. All authors read and approved the manuscript.

## DATA AVAILABILITY

Raw sequencing data, processed count matrices and brightfield images (HE-images) will be deposited and made publicly available upon publication.

## CODE AVAILABILITY

All code, data and results that relate to the content of this manuscript are available as a github repository found at https://github.com/almaan/her2st. The repository also includes results and all code used to generate these as well as the figures.

The presented results (clustering and single cell integration) can be interactively explored through a Shiny app, which is found at the aforementioned github repository, further instructions regarding how to open and orient this environment are provided at said location as well.

## METHODS

### Array production

The array production has already been described in previous publications.[18,50] Briefly, the microarrays were generated as a 33×35 grid of printed spots with a 200μm center-to-center distance of 100μm between each capture location (spot). A total of 1007 spots were printed with unique DNA oligonucleotides (spatial barcodes) attached to oligo(dT) capture probes.

### Sample acquisition

All tumors used for this analysis were immediately frozen in −80°C after surgery and trimming of fat, then stored in a tumor bank until the start of the experiment. For each tumor a different section - to that used in the ST experiments - was subjected to IHC and PAM50 analysis for classification of subtypes. All analyzed sections stained positive for HER2 and were classified as HER2 positive tumors by PAM50.

### Tissue handling, staining and imaging

These steps have previously been described in.[18] In short, fresh frozen material was sectioned at 16μm. After placing the tissue on top of the barcoded microarray, the glass slide was warmed at 37 °C for 1 min for tissue attachment and fixated in ∼ 4% NBF (neutral buffered formalin) for 10 min at room temperature (RT). The slide was then washed briefly with 1x PBS (phosphate buffered saline). The tissue was dried with isopropanol before staining. The tissue was stained with Mayer’s hematoxylin for 4 min, washed in Milli-Q water, incubated in bluing buffer for 2 min, washed in Milli-Q water, and further incubated for 1 min in 1:20 eosin solution in Tris-buffer (pH 6). The tissue sections were dried for 5 min at 37 °C and then mounted with 85% glycerol and a coverslip. Imaging was performed using the Metafer VSlide system at 20x magnification. The images were processed with the VSlide software (v1.0.0). After the imaging was complete, the cover slip and remaining glycerol were removed by dipping the whole slide in Milli-Q water followed by a brief wash in 80% ethanol and warming for 1 min at 37 °C.

### Permeabilization and cDNA synthesis

Permeabilization and cDNA synthesis were carried out as previously described, but with substitution of the Exonuclease I buffer pre-permeabilization treatment with a 20 min incubation at 37 °C in 14U of collagenase type I (Life Technologies, Paisley, UK).[18] The Exonuclease I buffer was diluted in 1x HBSS buffer (Thermo Fisher Scientific, Life Technologies, Paisley, UK) supplemented with 14μg BSA followed by an incubation in 0.1% pepsin-HCl (pH 1) for 10 min at 37 °C. A cDNA-mix containing Superscript III, RNaseOUT, DTT, dNTPs, BSA and Actinomycin D was added and the slide incubated at 42 °C overnight (∼18 h). The tissue was washed with 0.1x SSC between each incubation step.

### Tissue removal and cDNA release from the surface

Tissue removal, as well as the release of cDNAs from the surface have been described in prior publications.[18] In brief, beta-Mercaptoethanol was diluted in RNeasy lysis buffer and samples were incubated for 1 h at 56 °C. The wells were washed with 0.1x SSC followed by incubation with proteinase K, diluted in proteinase K digestion buffer, for 1 h at 56 °C. The slides were then washed in 2x SSC + 0.1% SDS, 0.2x SSC followed by 0.1x SSC and dried. The release mix consisted of second strand buffer, dNTPs, BSA and USER enzyme and was carried out for 2h at 37 °C. After probe release, the 1007 spatial spots containing non-released DNA oligonucleotide fragments were detected by hybridization and imaging, in order to obtain Cy3-images for image alignment and spot detection, as described previously.[18]

### Library preparation and sequencing

The protocol followed the same preparation procedures as described earlier in [18], but were carried out using an automated pipetting system (MBS Magnatrix Workstation), also previously reported.[51] In general, second strand synthesis and blunting were carried out by adding DNA polymerase I, RNase H and T4 DNA polymerase. The libraries were purified and amplified RNA (aRNA) was generated by a 14h *in vitro* transcription (IVT) reaction using T7 RNA polymerase, supplemented with NTPs and SUPERaseIN. The material was purified and an adapter ligated to the 3’-end using a truncated RNA ligase 2. Generation of cDNA was carried out at 50 °C for 1 h by Superscript III, supplemented with a primer, RNaseOUT, DTT and dNTPs. Double stranded cDNA was purified, and full Illumina sequencing adapters and indexes were added by PCR using 2xKAPA HotStart ready-mix. The number of amplification cycles needed for each section was determined by qPCR with the addition of EVA Green. Final libraries were purified and validated using an Agilent Bioanalyzer and Qubit before sequencing on the NextSeq500 (v2) at a depth of ∼100 million paired-end reads per tissue section. The forward read contained 31 nucleotides and the reverse read 46 nucleotides.

### Mapping, gene counting and demultiplexing

These steps were carried out in a similar fashion to what previously have been described.[18] The forward read contained the spatial barcode and a semi-randomized UMI sequence (WSNNWSNNV, with: W - A/T, S - G/C, N - A/C/T/G and V - A/C/G) while the reverse read contained the transcript information and was used for mapping to the reference GRCh38 human genome. Before mapping the reads with STAR [52], the reverse reads were first quality trimmed based on the Burrows-Wheeler aligner, long homopolymer stretches were also removed. Multi-mapped reads, i.e. reads mapping to multiple loci in the genome, were discarded after mapping with STAR. HTSeq-count with the setting *-intersection-nonempty*, was used to generate gene counts, using an Ensembl reference file (v. 86).[53] The remaining reads were provided as input to TagGD demultiplexing using the 18 nucleotides spatial barcode.[54] The demultiplexed reads were then filtered for amplification duplicates using the UMI with a minimal hamming distance of 2. The UMI-filtered counts were used in the analysis. The analysis pipeline (1.6.0) is available at https://github.com/SpatialTranscriptomicsResearch/st_pipeline.

### Pre-processing

Raw data was merged from 6 section gene count matrices for samples A, B, C and D and 3 section gene count matrices for samples E, F, G and H. The merged expression matrices were enriched for genes matching the biotypes protein_coding, IG_C_gene, IG_J_gene, IG_V_gene, TR_C_gene, TR_J_gene and TR_V_gene. In addition, each merged expression matrix was filtered from ribosomal protein genes (RPL and RPS), mitochondrial genes (MT-) and MTRNR genes as well as genes expressed in fewer than 10 spots across the whole merged dataset. Spots with fewer than 300 unique features (genes) were also removed from the merged datasets.

### Normalization and feature selection

The merged data was first normalized using the regularized negative binomial regression method implemented in the *SCTransform* function from Seurat (v3.1.4) R package.[55] The number of variable genes selected with *SCTransform* was determined by applying a residual variance cutoff of 1.1 (variable.features.rv.th = 1.1) with the additional parameter settings; return.only.var.genes = FALSE and variable.features.n = NULL. In the subsequent patient-based analysis, we applied the same normalization scheme but with an additional batch correction term to adjust for technical differences across replicate tissue sections (vars.to.regress = section).

### Dimensionality reduction

Before running dimensionality reduction, the set of highly variable genes as defined by the *SCTransform* method was reduced to a smaller set of genes as described below. First, we hypothesized that the most relevant features should not only have high variance, but also show positive spatial autocorrelation. We therefore devised a method to rank the variable features by spatial autocorrelation by computing the pearson correlation coefficient for each gene between the expression vector and the spatial lag vector (defined as the summed expression in the adjacent neighboring spots over all spots). Variable genes with a correlation coefficient larger than 0.1 were therefore kept in the reduced gene set. We also identified 21 highly variable genes which contributed to form a ring like pattern in several capture areas (Supplementary Data 4). This effect was not found in all biological replicates from the same tissue biopsies and was therefore concluded to be a source of technical variation. All 21 genes were excluded from the reduced gene set. For each patient dataset, the reduced set of highly variable and spatially correlated genes was used as input for a Non-negative Matrix Factorization (NMF) computation with 10 factors using the *RunNMF* function from the STUtility R package.[56] Each factor was then visualized as a spatial heatmap colored by factor value and factors with consistent patterns across replicate tissue sections were kept for subsequent analysis steps.

### Expression-based clustering

First, a Shared Nearest Neighbor (SNN) graph was constructed from the selected NMF factor matrix with the *FindNeighbors* function in Seurat. This SNN graph was then used to identify clusters of spots using the modularity based clustering algorithm implemented in the *FindClusters* function in Seurat. The resolution parameter was set to 0.4 for all samples.

### Marker detection

For each patient dataset, a Wilcoxon signed-rank test was performed using the *FindAllMarkers* function in Seurat to find differentially expressed genes within each cluster. The function performs the test pairwise between each cluster and its background (all other spots in the dataset). The resulting table of gene markers was filtered to include genes with an adjusted p-value lower than 0.01 and an average log fold (natural logarithm) change higher than 0.15, thus omitting down-regulated genes.

### Cluster annotation

Each set of differentially up-regulated genes were subjected to enrichment analysis using the Gene Ontology – Biological Processes (GO:BP) database and the enricher function from the g:profiler R package with an adjusted p-value cutoff of 0.05. Each cluster was then manually annotated using the top enriched pathways and up-regulated marker genes as basis.

### Cluster overlap between patients and core signature extraction

To check for overlapping gene signatures between clusters from different patients, we computed the Jaccard index for all pairs of cluster gene sets (up-regulated DE genes). These values were first used to compute a distance matrix (euclidean distance) from which a dendrogram was constructed (R package *hclust*) with the agglomeration method set to “complete”. This dendrogram was then cut into 5 groups using the *cutree* function (R package *stats*) with k (number of clusters) set to 5. Then, for each group of clusters we extracted all genes that were shared between at least two clusters. For two of the groups, zero genes were shared across all clusters and these groups were excluded. For the remaining 3 groups, we defined a core signature as the genes that were shared across at least 80% of the clusters.

### Single Cell Data

We downloaded the single cell data related to the publication [30]. Only cells originating from the HER2-positive patients were used in our analysis. We used the same labels as in the figures of the single-cell resource, with the exception of Plasmablasts which we here refer to as Plasma Cells.

### Spatial Mapping of Single Cell Data

To infer the spatial organization of certain cell types we used a method developed to integrate spatial and single cell data, implemented and available as a python package (*stereoscope*, v.0.2, https://github.com/almaan/stereoscope). The method is based on a probabilistic model which assumes that both single cell and spatial RNA-seq data follow a negative binomial distribution. By using annotated single cell data in combination with spatial transcriptomics data it estimates proportions of every cell type (present in the single cell data) at each spatial capture location.[22]

To conduct the spatial mapping of cell types, we only included cells from the five HER2-positive patients found in the single cell data, all ST-sections were used. In total three analyzes were conducted, with the only difference being the labels used for the single cell data. As mentioned in the main text, every cell was assigned to a type within each of the three tiers *major, minor* and *subset*. For respective tier, we subsampled the single cell data set, according to the following scheme: (i) If a type had fewer than 25 members, exclude the type; (ii) if a type had more than 25 members but less than or equal to 500 members, include all cells; (iii) if a type had more than 500 members, randomly select 500 of these. Next, the subsampled sets were spatially mapped, one by one, onto the ST data.

A custom gene list of 5540 members, representing the union of the 5000 (highest expressed) genes in the single cell data and cell type marker genes, were used for the proportion inference, see Supplementary Data 11. 50000 epochs and a batch size of 2048 were used for all tiers, in both steps of the *stereoscope* procedure. Default values were used for all remaining parameters.

### Cell type co-localization

We use spotwise Pearson correlation between the estimated cell type proportion values as a proxy for cell type co-localization; with high positive correlation being indicative of types that exhibit similar spatial distributions and the opposite being true for negative values. To estimate the confidence interval for each of the correlation values, we used a bootstrap approach. The Pearson correlation was computed for each of 10000 bootstrap samples (sampling from all spots with replacement), forming a distribution of correlation values for each pair of types. The mean of each distribution was taken as a representative correlation value, and a 95% confidence interval was defined by the 2.5th and 97.5th percentiles. Pairs where the confidence interval included zero were considered as not statistically significant, indicated with a gray border.

### Region based enrichment/depletion of cell types

The enrichment, or alternatively depletion, of the mapped cell types in relation to spatial regions (e.g., manual annotations or clusters) were assessed by the following procedure: First, the average proportion value was computed within each of the regions, referred to as the *true average*. Next, we permuted the spot indices for the proportion estimate vectors 10000 times, while maintaining the original indices for the annotated regions. In other words, the proportion estimates were shuffled w.r.t to their spatial location. Average proportion values of the annotated regions were determined for each permutation, constituting the set of *permuted averages*. We then computed the differences between the true average and the all permuted averages. Finally, the mean value of the differences divided by the standard deviation of these differences were taken as the *enrichment score* for the respective regions.

Upon visualization, the *enrichment score* of a type within a certain region is represented by two features, the marker size and it’s color. We let the marker size be proportional to the absolute value of the enrichment score, while the color indicates the sign (red for negative and green for positive). To summarize, red markers are indicative of a type being depleted in a certain region, green markers of enrichment; the larger the marker, the larger the effect.

### TLS Signatures

The method we devised to obtain a TLS signature can be decomposed into two steps: (1) associating a TLS-score to each spatial location and (2) modelling the contribution of each gene to this score. We base the TLS-score in the proportion values estimated for the major cell type tier; first a *raw TLS-scores* are computed, taken as the product between B and T-cell proportions multiplied by the scalar 2. In theory — assuming unbiased and independent sampling from a large population of cells with the same type composition as the spot — this represents the probability that two randomly selected cells is a B respectively T-cell. The raw TLS-score is then adjusted by subtracting the average probability of picking any cell type pair in the associated spot, this is the final TLS-score used in the subsequent steps.

In the second step, we consider the (adjusted) TLS-score at a given spot as a function of its gene expression. The gene expression values are normalized accordingly: First, all elements of a spot’s expression vector are divided by its library size (the sum of all elements); second, the expression vector associated to each gene is divided by its standard deviation (computed after the preceding library size division). Letting **y** represent the S-dimensional (S being the number of spots) TLS-score vector, **X** the SxG (G being the number of genes) normalized expression vector, **β** a G-dimensional vector of coefficients and β_0_ a scalar representing the intercept value; we estimate values of **β** and β_0_ that minimize the loss function L(**β)** = ||**y -** (**Xβ +** β_0_**1**)**||^2^,** where **1** is a G-dimensional vector with all elements being 1. Implementation wise, we used the *OLS class* from the *linear_model* module in the python package statsmodels (version 0.11.0) for the purpose of finding the least square estimates.

The genes qualifying as members in the final TLS-signature is determined by ordering the coefficient values from largest to smallest, considering the values as a function (*f)* of the gene’s rank. The resulting curve is then smoothed with a gaussian filter, and the second order differences of this smoothed curve are computed, representing an approximation of *f*’s second derivative. Using the same gaussian filter as previously mentioned, the second derivative approximation is smoothed. The gene coefficient for which the smoothed second derivative approximation obtains a value below zero is taken as the lower bound (threshold), hence all genes with a coefficient having a lower rank than this will be excluded. The gaussian filtering was performed by using the *gaussian_filter* function from scipy’s (version 1.4.1) *ndimage* module; the sigma parameter was set to 10 whilst default values were used for the remaining parameters. Applying the aforementioned procedure to all replicates of patient G and H, we obtained a signature of 171 genes, full list in Supplementary Table 2.

Functional enrichment of the gene signature was performed by using the *g:Profiler* python package (version 1.0.0), we queried against GO:BP (GO Biological Processes) and selected all terms that were significantly enriched (having an adjusted p-value smaller than 0.05). The complete set of these pathways are found in Supplementary Data 12.

### Spot-wise pathway enrichment

To assess enrichment of a given gene set (here associated to a given functional pathway) spatially, we used the following approach. Let G be all genes present in the spatial data, and let Q_P_ be the set of all genes associated with the pathway P for which enrichment should be examined. Also, for each gene subtract its average expression, then within each spot rank the genes according to their adjusted expression levels (from highest to lowest). Let T_n_(s) be the set of *n* highest ranked genes within spot *s.* Now, for each spot construct a 2×2 contingency table of the following character:

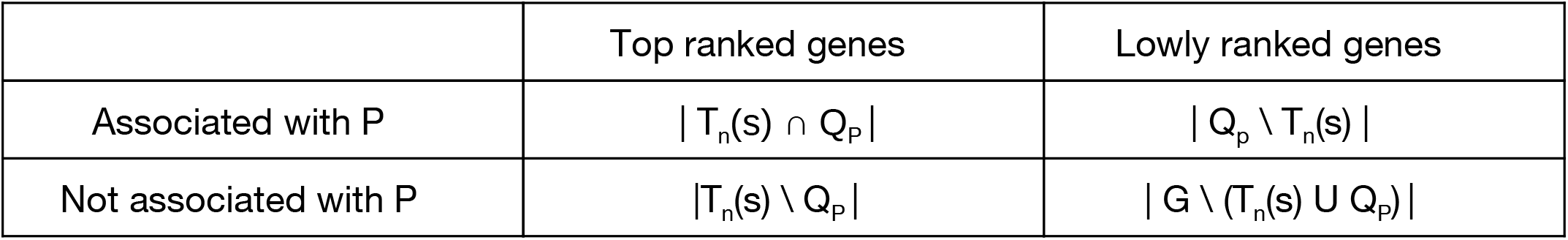

Then conduct a Fisher’s exact test, to calculate the probability (p(s)) of observing this partitioning of genes among the two variables, assuming that the genes associated with P are equally distributed over the top (T_n_(s)) and lower ranked genes. The enrichment (E_P_(S)) of P for spot *s* is then taken as : E_p_(s) = −log2(p(s)). These are the values visualized in Figure 4E.

## Supplementary

**Supplementary Figure 1.**
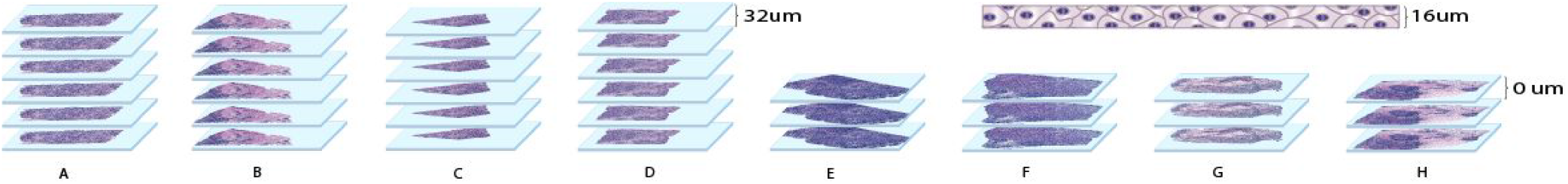
Experimental setup. Six cryosections were taken with a distance of 32 um from patient A-D respectively and three consecutively cut cryosections were taken from patient E-H respectively. Each section was taken with a thickness of 16 um and placed on a ST-array.

**Supplementary Figure 2.**
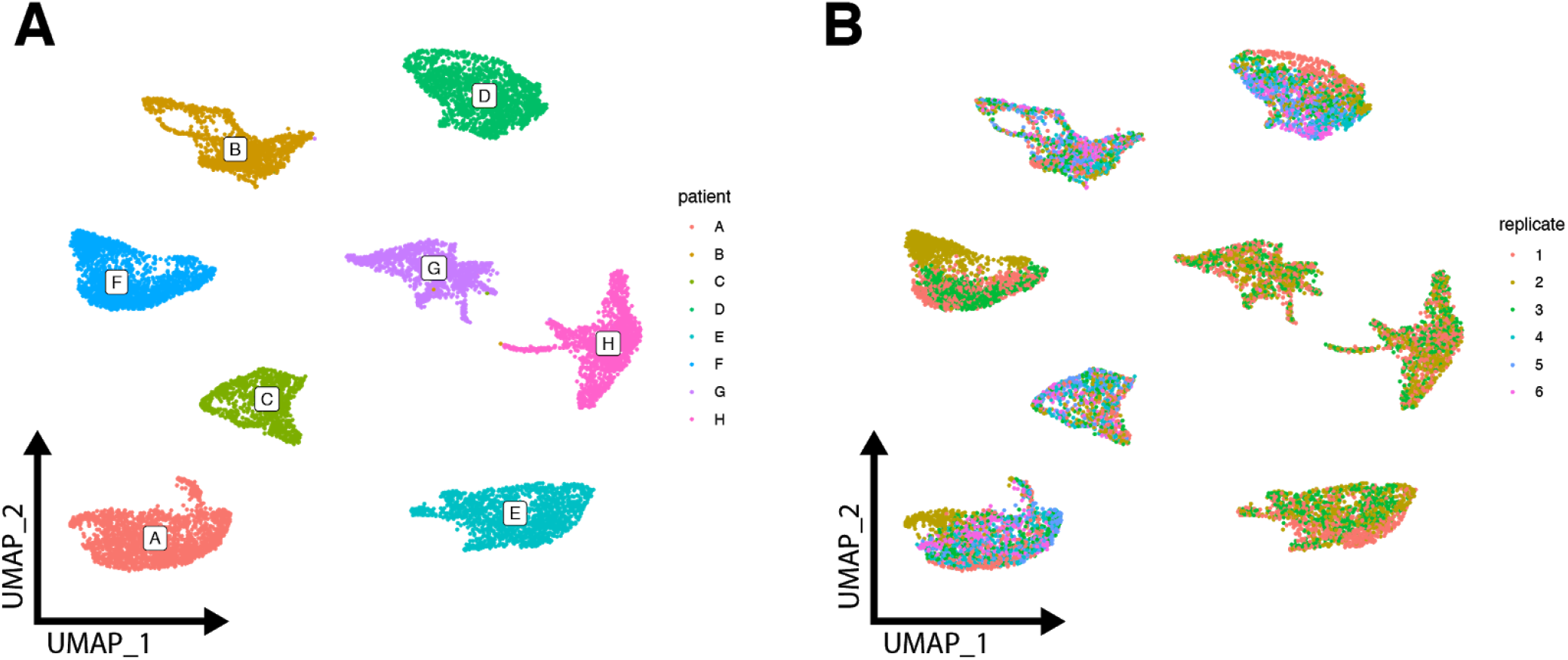
UMAP-plot. **A.** UMAP visualization of all spots colored by patient. **B.** UMAP visualization of all spots colored by replicate.

**Supplementary Figure 3.**
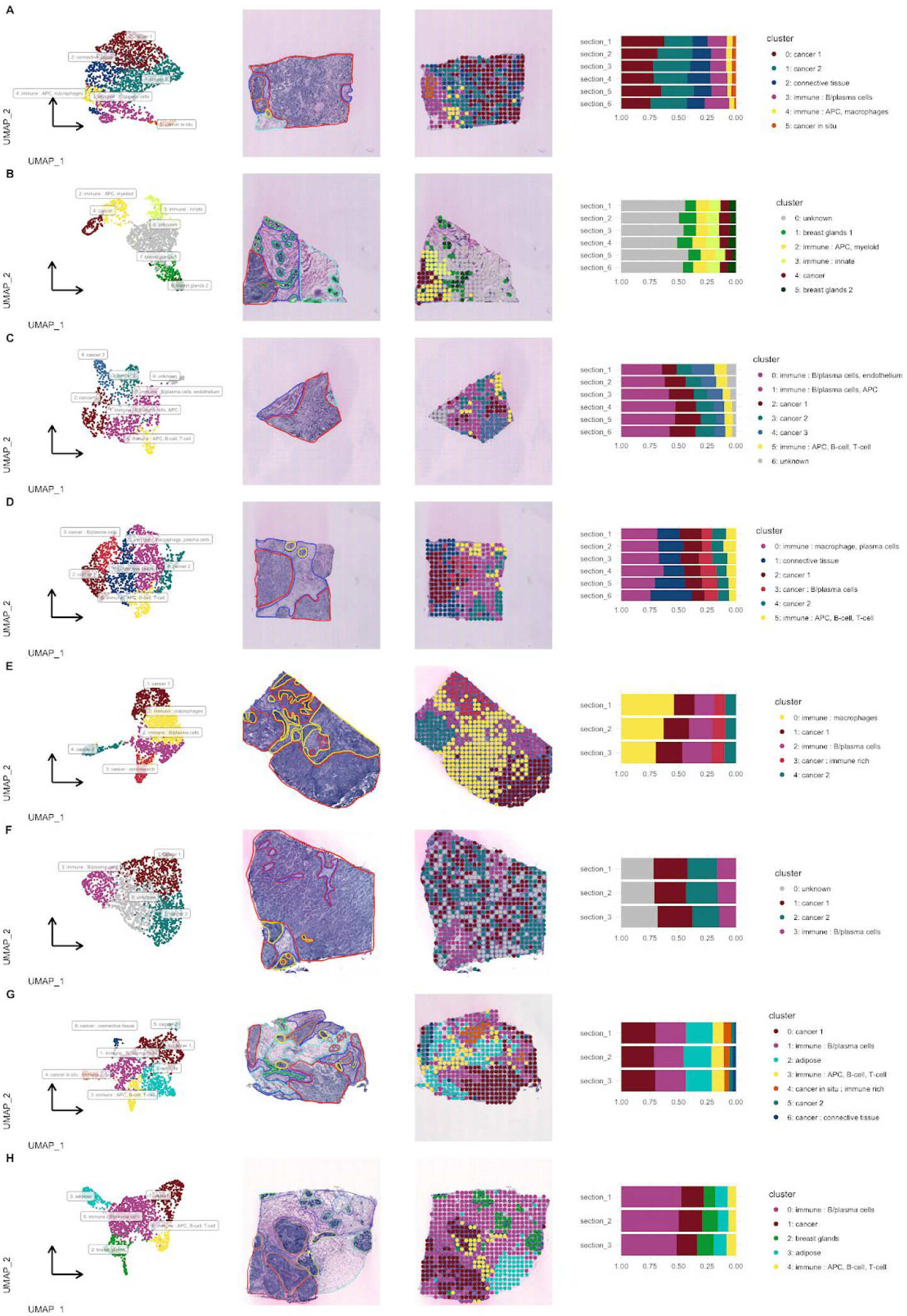
Expression-based clustering overview. **Column 1.** UMAP visualization of spots colored by cluster. **Column 2**. Annotations made by a trained pathologist into six distinct categories: Invasive Cancer (red), Adipose tissue (cyan), Connective tissue (blue), Breast glands (green), *in situ* cancer (orange) and Inflammatory cells (yellow). **Column 3.** Spatial visualization of clusters. **Column 4.** Proportions of spots belonging to each cluster across replicate sections.

**Supplementary Figure 4.**
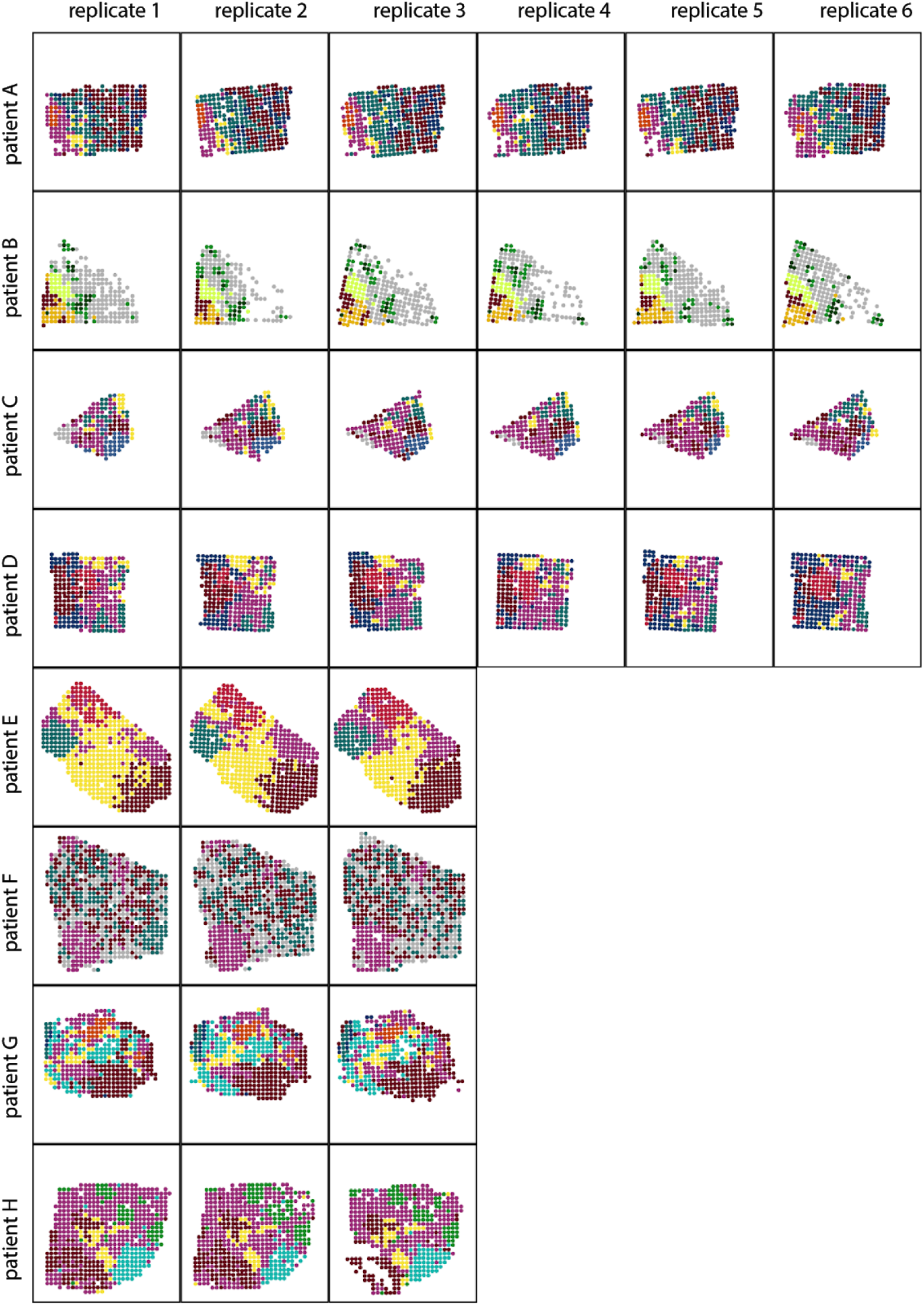
Spatial arrangement of all expression-based clusters. Spatial visualization of clusters across all replicate tissue sections for each patient (A-H). Spots with the same colors belong to the same clusters. Clusters are not shared between patients.

**Supplementary Figure 5.**
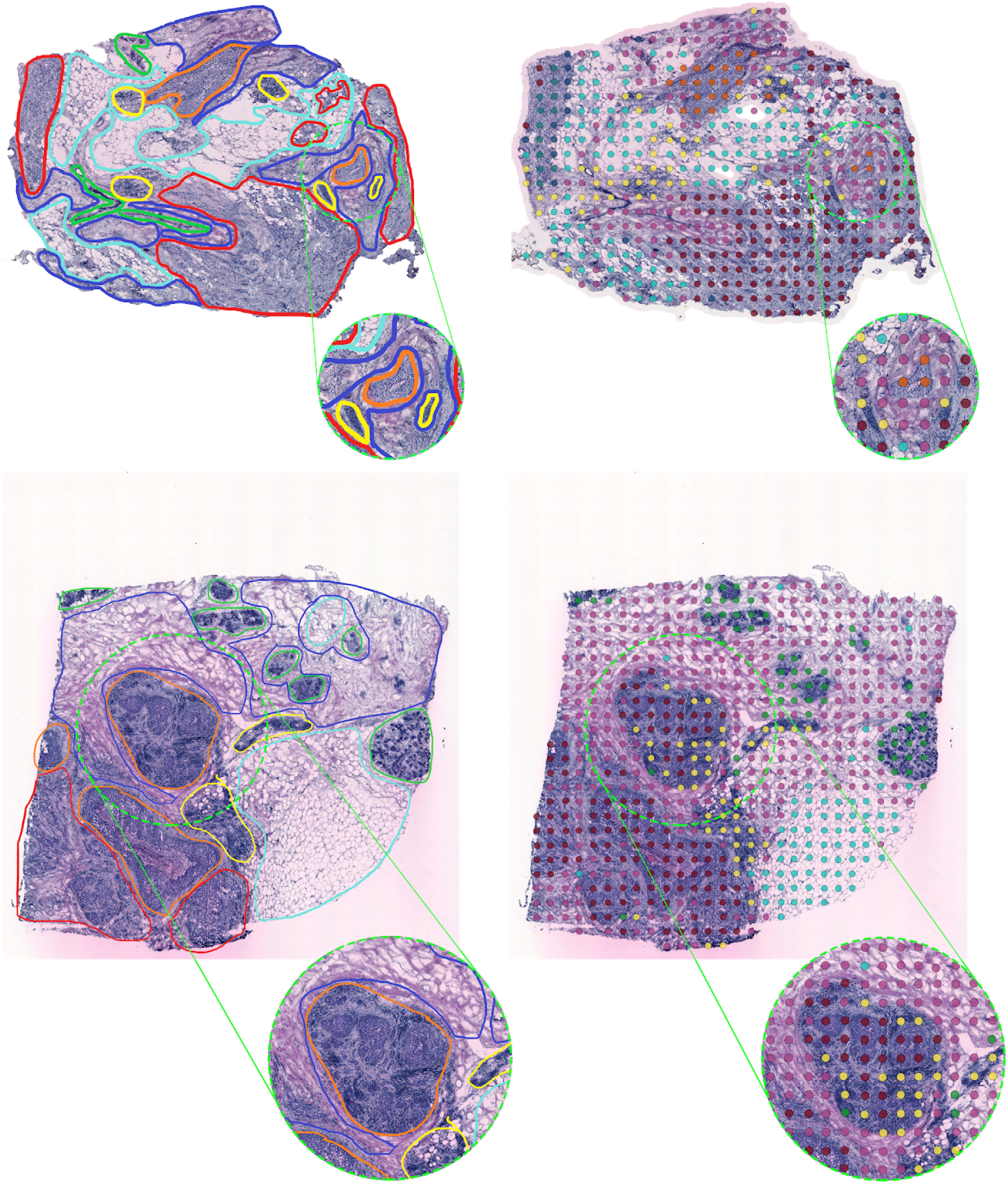
Patient G and H Zoom-in of specific regions of interest. (Top row) A region in patient G consisting of only three spots and aligning with the *in situ* cancer regions is assigned to a different cluster than its spatial neighbors (orange), the same cluster as the rest of the *in situ* cancer spots are found in. (Bottom row) A region with *in situ* region in patient H is populated by two different clusters (purple and yellow)

**Supplementary Figure 6.**
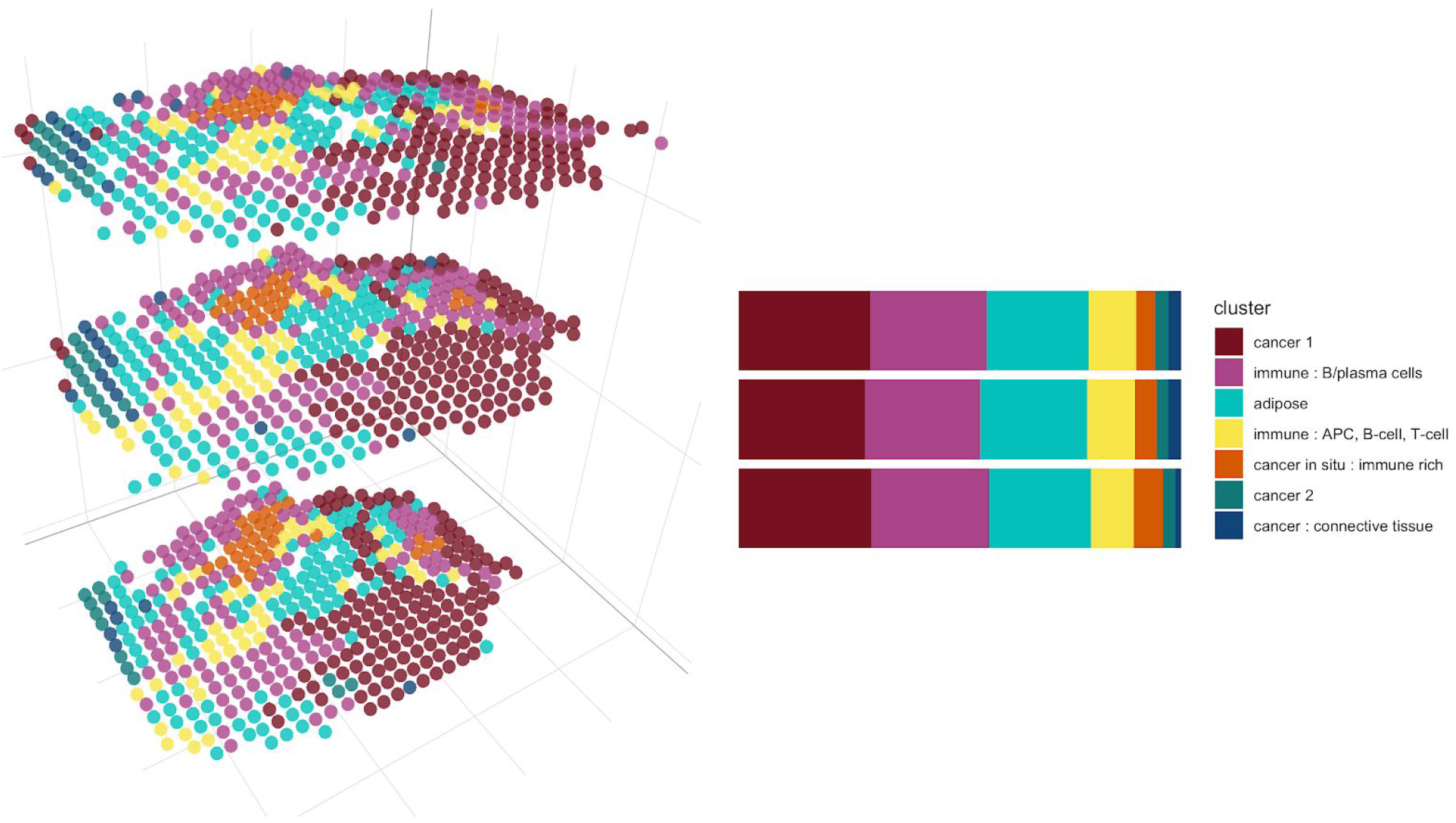
3D visualization of Patient G. The three replicates taken from Patient G, visualizing the expression based clusters’ distribution in all three dimensions. Distances in the XY-plane and Z-axis are, for ease of visualization, not depicted in the same scale.

**Supplementary Figure 7.**
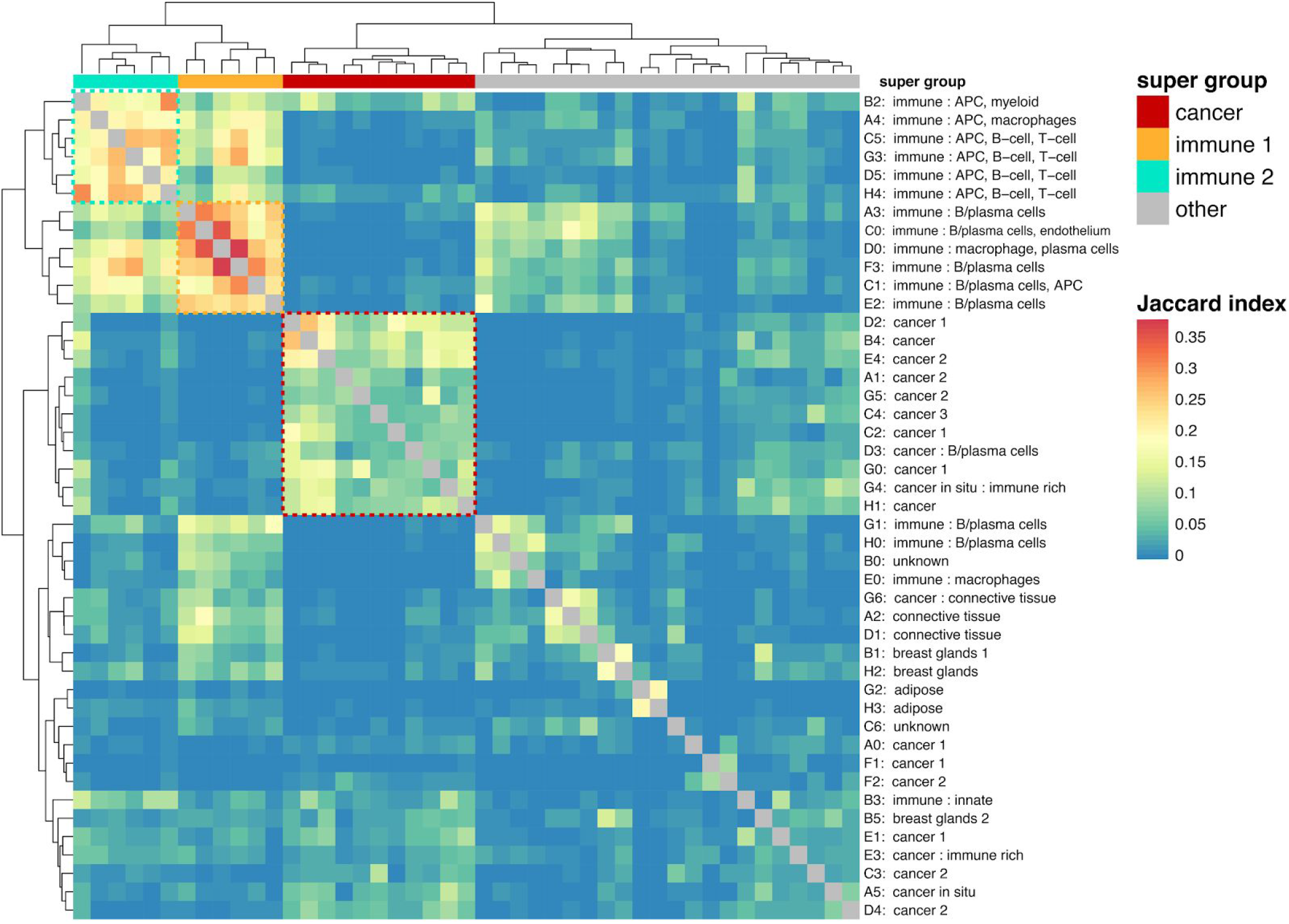
Clusters-of-clusters plot. Heatmap of Jaccard indices calculated across cluster geneset paris. The three cluster supergroups are highlighted by the dashed boxes, each defined by their upregulation of core signature genes. Group 1: cancer, group 2 and 3: immune related.

**Supplementary Table 1.** Receptor Status. ER and PgR receptor status for all patients (A-H) used during the tumor classification. Only Patient B (bold) has positive PgR status.

| Patient | ER | PgR |
| --- | --- | --- |
| A | - | - |
| **B** | - | **+** |
| C | - | - |
| D | - | - |
| E | - | - |
| F | - | - |
| G | - | - |
| H | - | - |

**Supplementary Figure 8.**
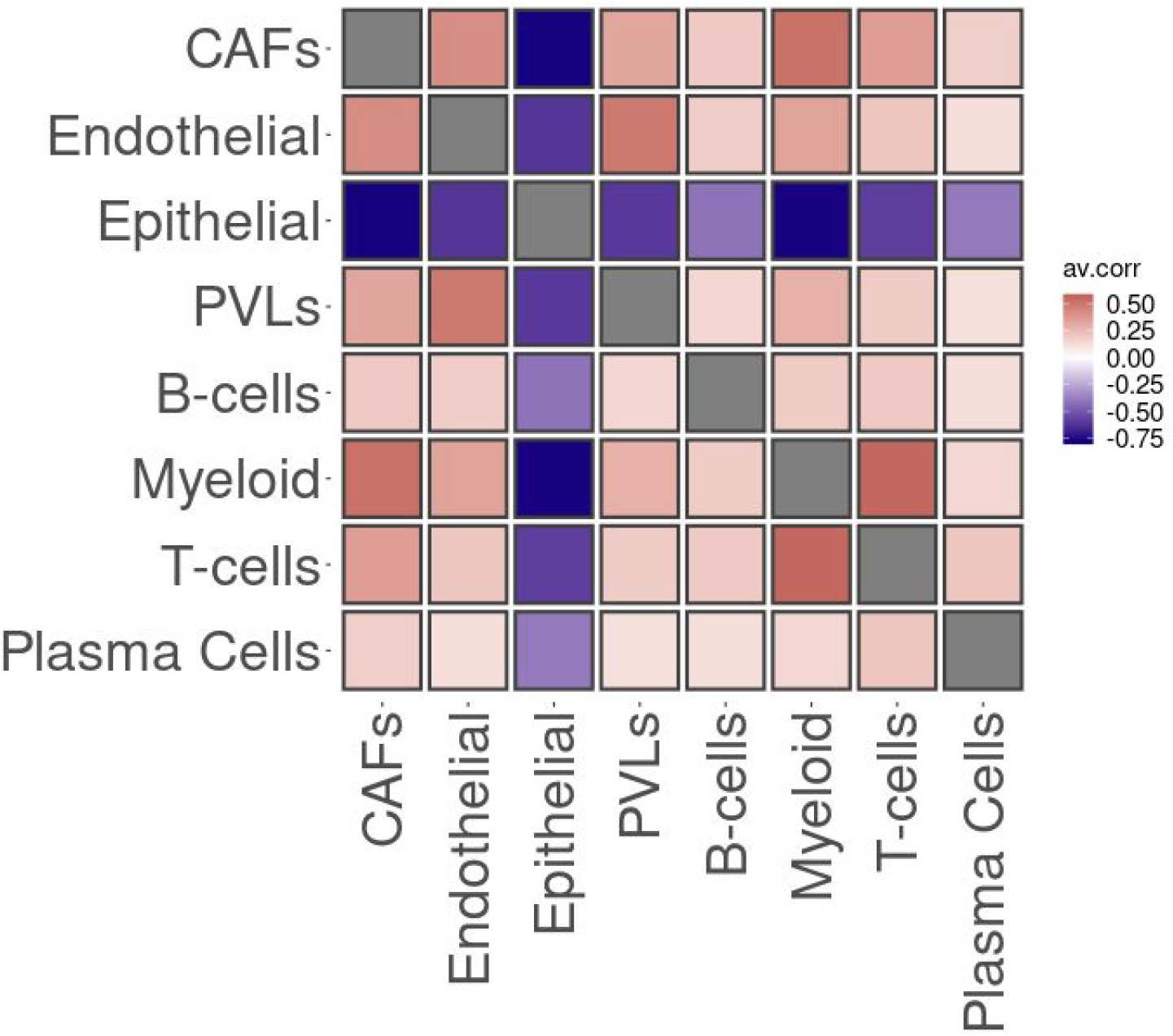
Correlation plot - Patient A, major tier. Correlation (Pearson) plot of cell type proportions across the spots, red is indicative of high spatial co-localization, blue is indicative of low spatial co-localization. The correlation values are computed over all six replicates of patient A, for the major tier.

**Supplementary Figure 9.**
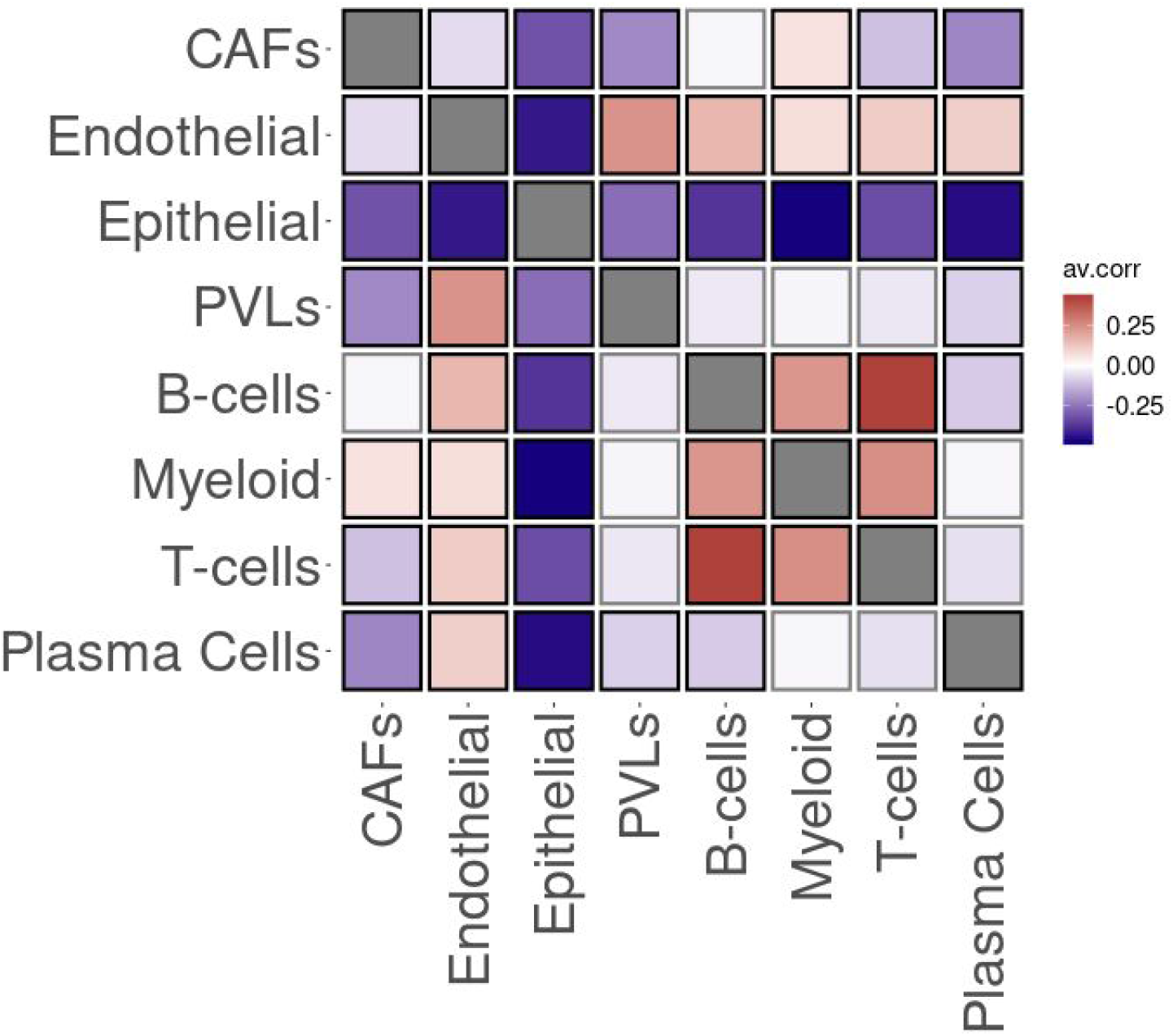
Correlation plot - patient G, major tier. Correlation (Pearson) plot of cell type proportions across the spots, red is indicative of high spatial co-localization, blue is indicative of low spatial co-localization. The correlation values are computed over all three replicates of patient G, for the major tier

**Supplementary Figure 10.**
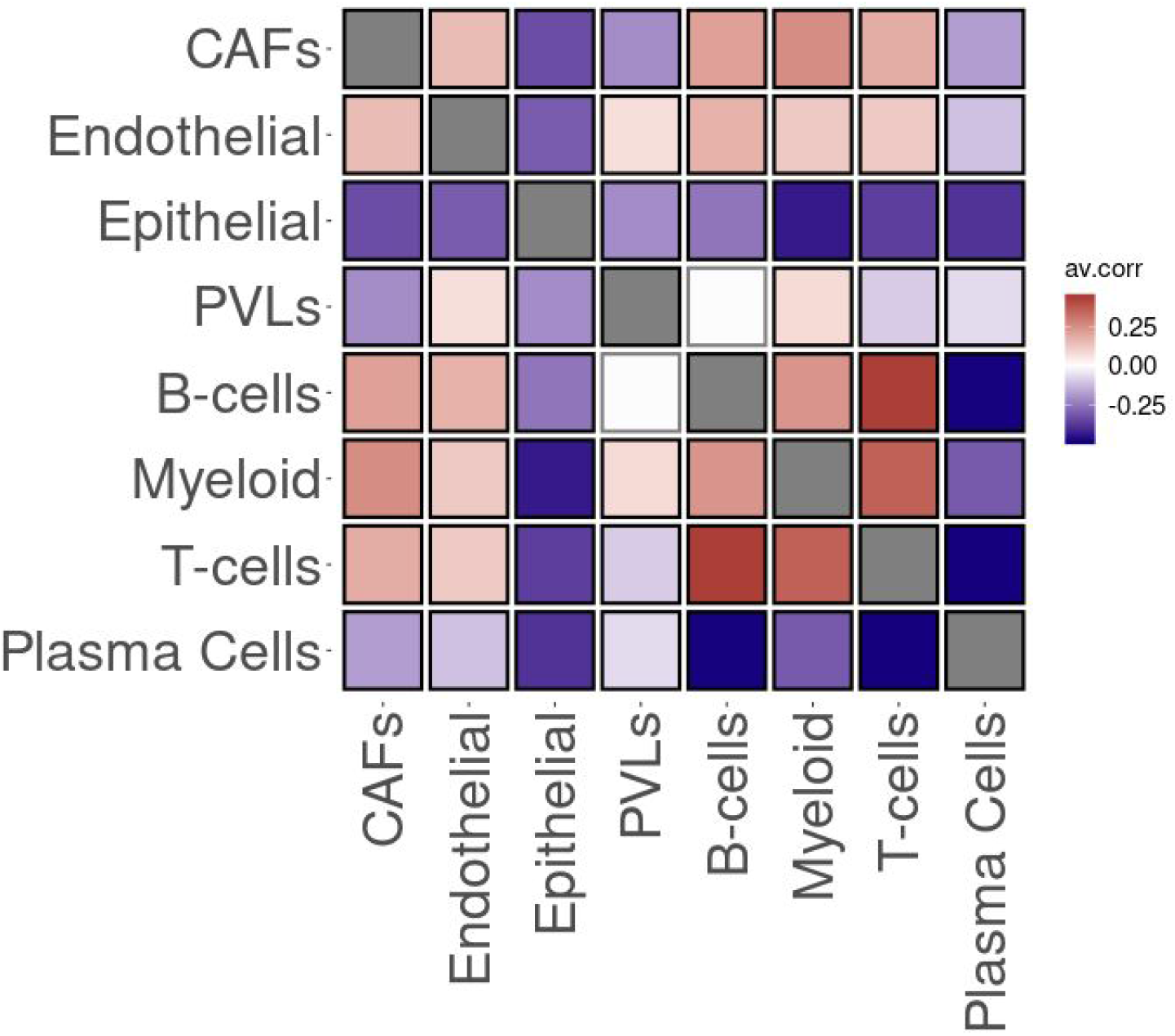
Correlation plot - Patient H, major tier. Correlation (Pearson) plot of cell type proportions across the spots, red is indicative of high spatial co-localization, blue is indicative of low spatial co-localization. The correlation values are computed over all three replicates of patient H, for the major tier.

**Supplementary Figure 11.**
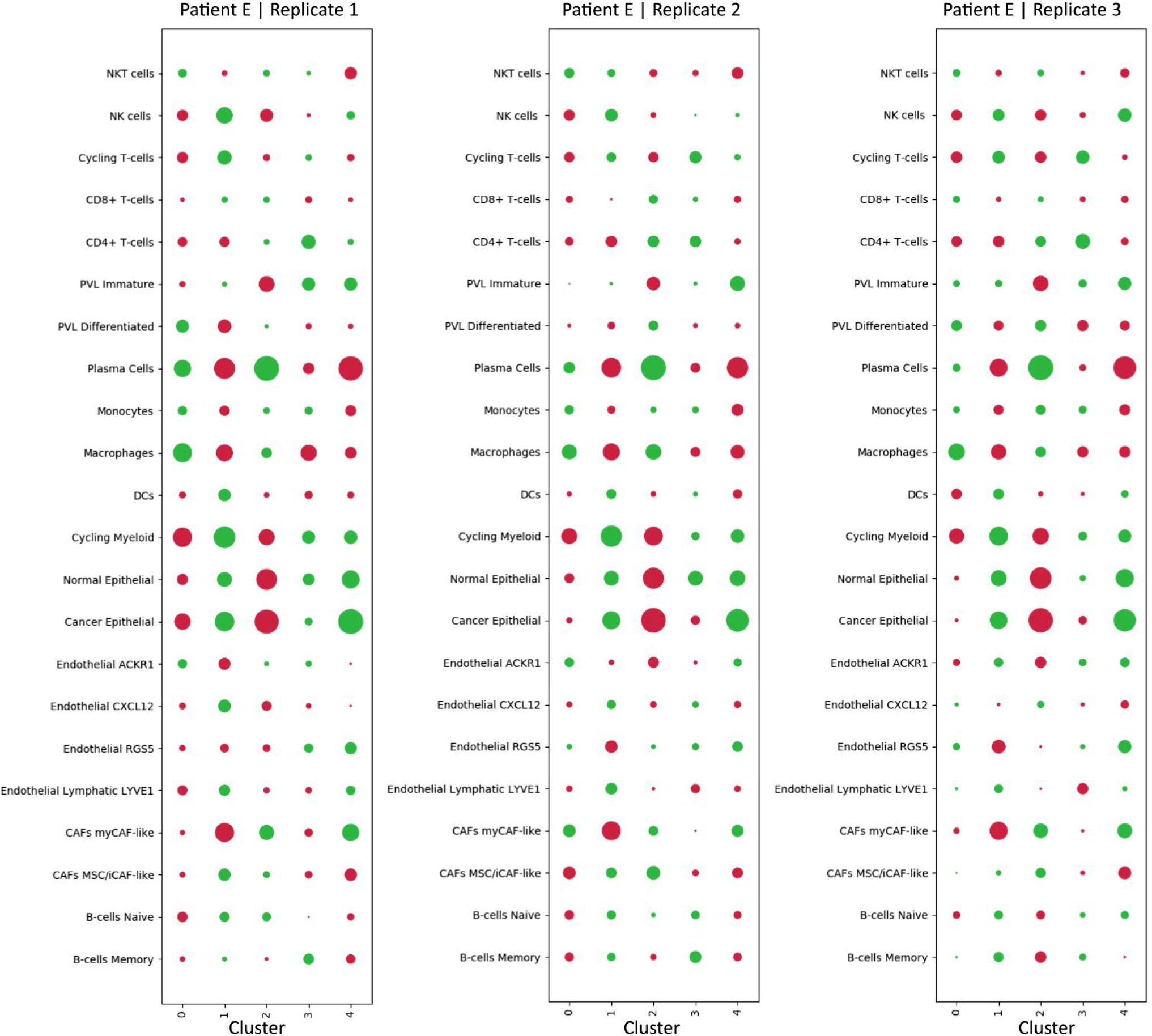
Enrichment in expression-based clusters, Patient E, minor tier. Patient E, enrichment of minor tier cell types within expression based clusters. Red is indicative of depletion, green of enrichment. Markersize is indicative of the extent of the effect (depletion or enrichment).

**Supplementary Figure 12.**
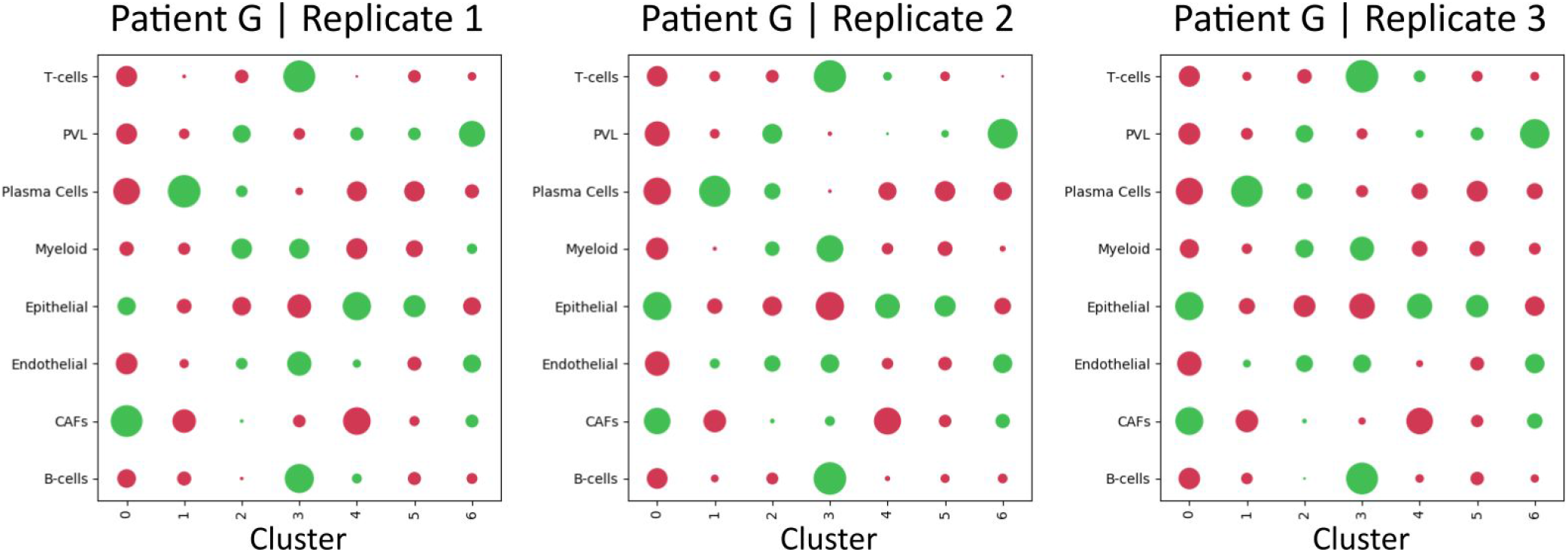
Enrichment in expression-based clusters, Patient G, major tier. Patient G, enrichment of major tier cell types within expression based clusters. Red is indicative of depletion, green of enrichment. Markersize is indicative of the extent of the effect (depletion or enrichment).

**Supplementary Figure 13.**
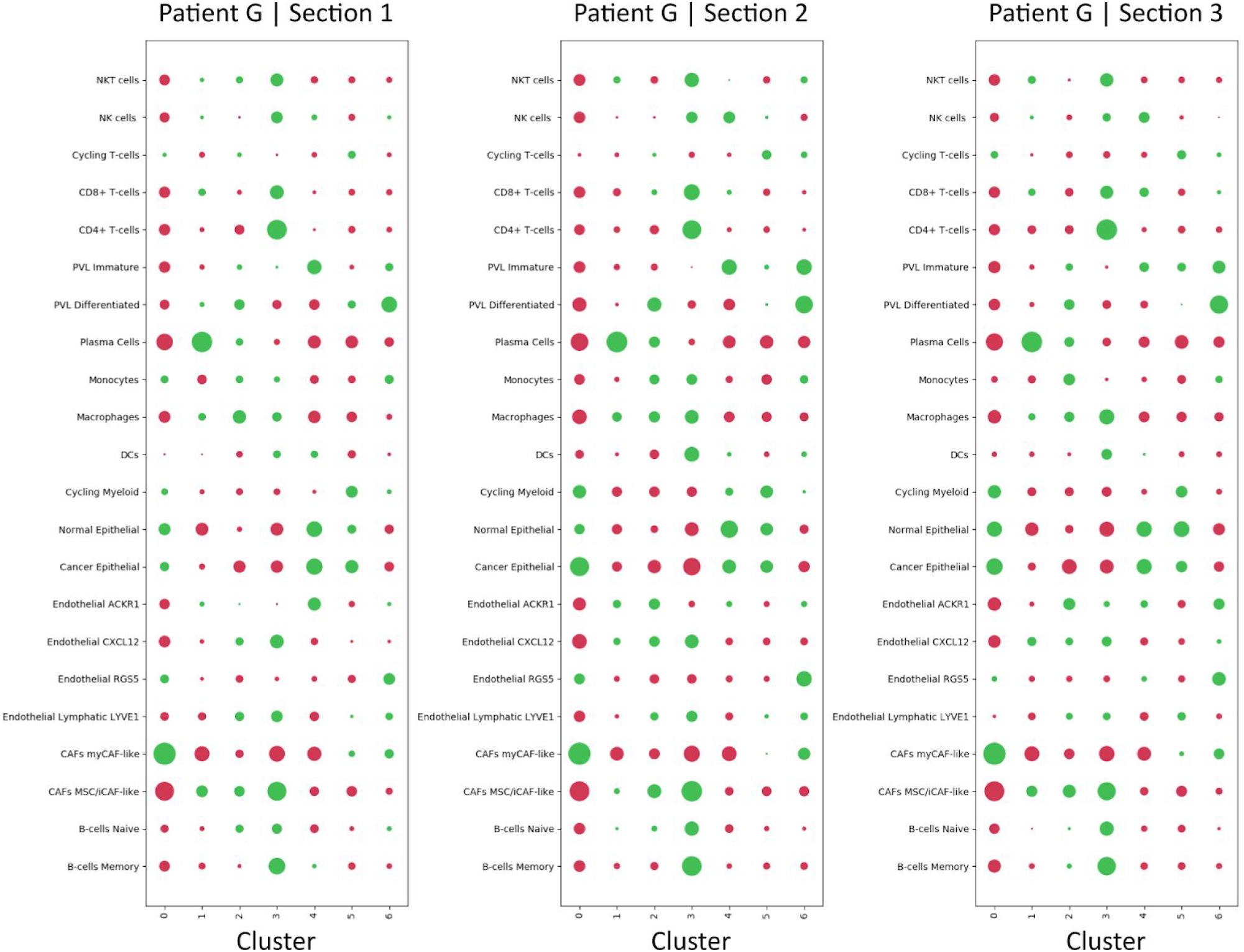
Enrichment in expression-based clusters, Patient G, minor tier. Patient G, enrichment of minor tier cell types within expression based clusters. Red is indicative of depletion, green of enrichment. Markersize is indicative of the extent of the effect (depletion or enrichment).

**Supplementary Figure 14.**
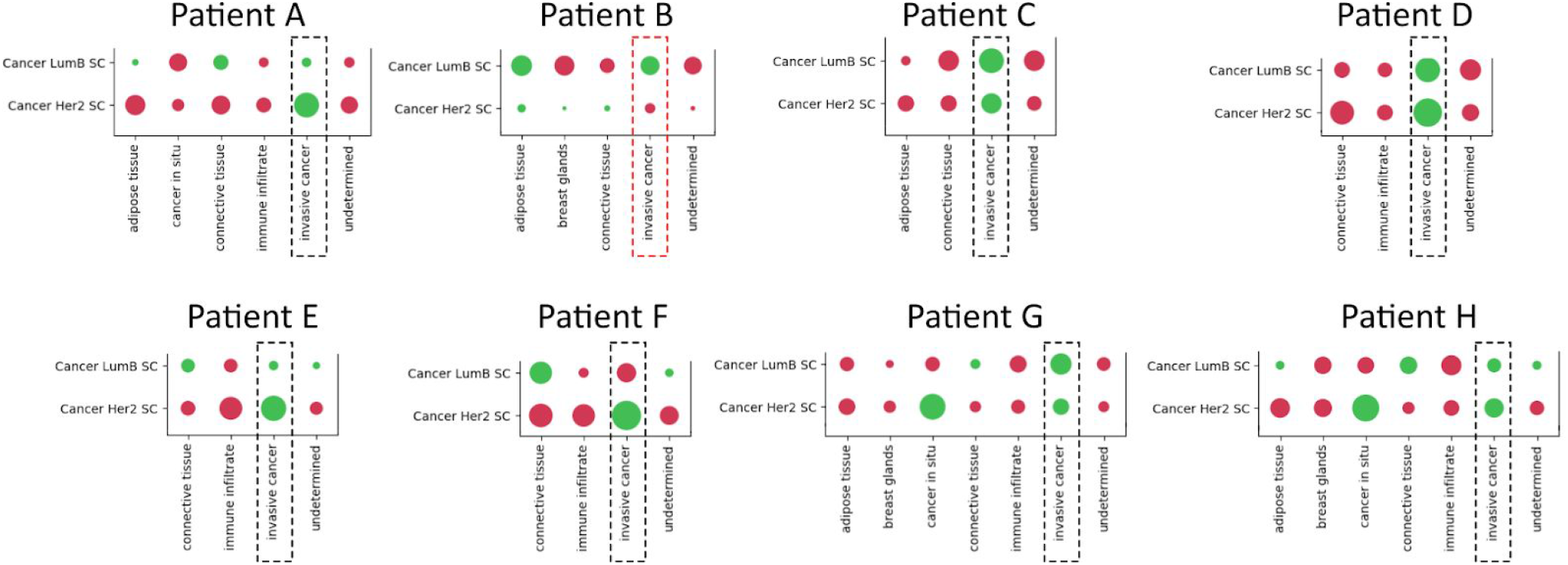
Cancer cell type enrichment across all patients. Enrichment/depletion plots of Her2 and LumB associated cancer cell types in respective patients in the regions defined by the pathologist. Dashed boxes indicate regions of invasive cancer. Patient B (red dashed box) is the only patient that exhibits depletion (red) of HER2 but enrichment (green) of LumB associated cancer cell types.

**Supplementary Section 1 | Prediction on External Data**

Having established good performance for our predictive model across techniques when applied to breast cancer data we wanted to assess its behavior when presented with other types of tissue. Thus we predicted TLS-score in samples from three different tissues: rheumatoid arthritis (RA), melanoma and developmental heart, see Supplementary Figure 15. Regions with high TLS-score in RA tissue overlapped with the areas annotated as immune infiltrates in the original publication, where they also discuss the likely presence of ectopic lymphoid structures (in this context, synonymous to TLS).[57] A similar trend was observed in the melanoma sample, where high TLS-scores were predicted in regions annotated as immune infiltrates.[58] In the healthy developmental heart, no signals above zero were observed for the predicted TLS-score, suggesting absence of TLSs, as expected in this tissue.[59,60]

**Supplementary Figure 15.**
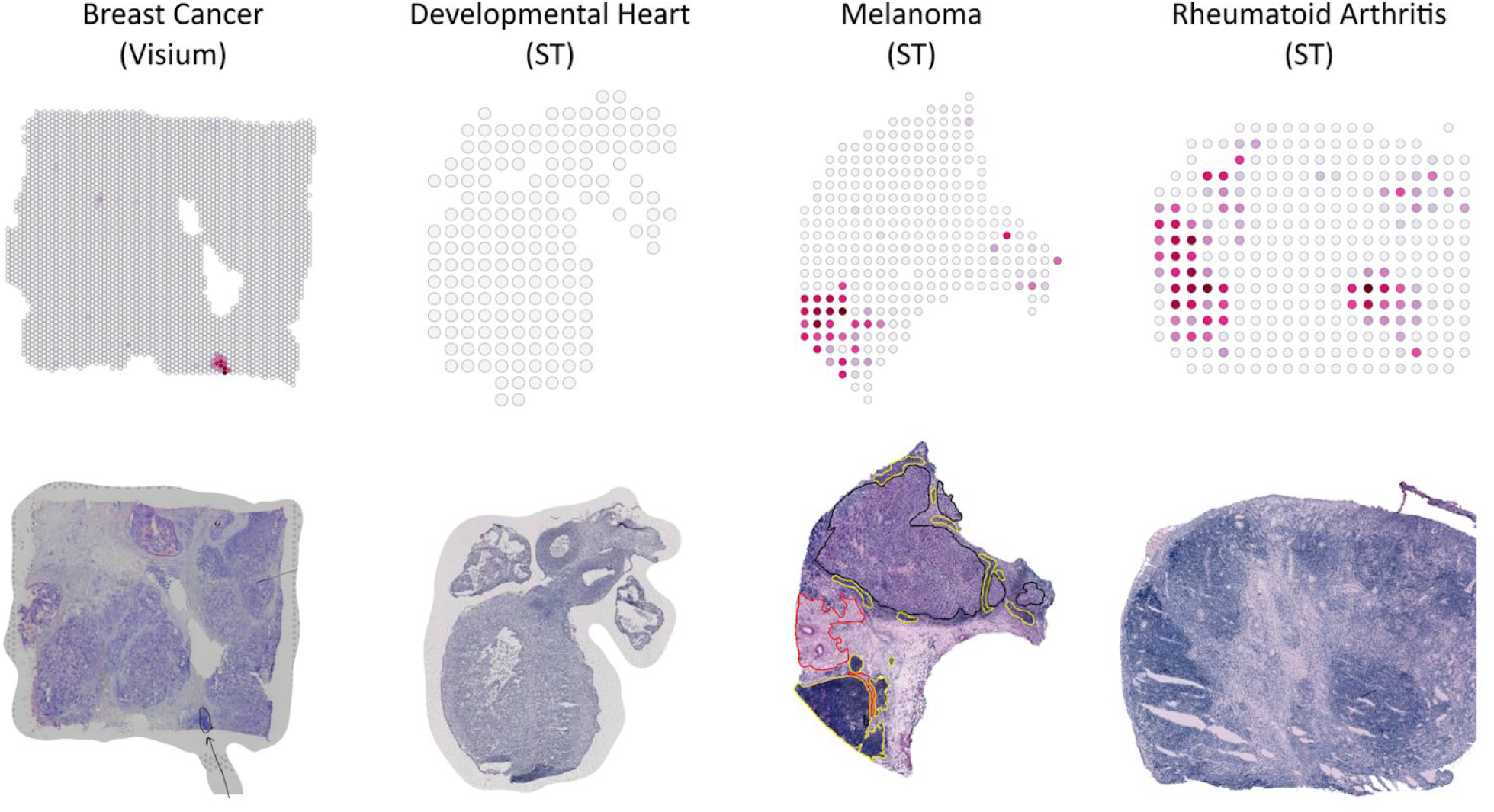
Validation on external data sets. (Top row) Predicted TLS-score of tissues from different platforms (Visium and ST) and tissues (breast cancer, developmental heart, melanoma and rheumatoid arthritis). (Bottom row) HE-images of the corresponding tissue. Our pathologist has annotated the Visium sample, with an arrow indicating a likely TLS site (the only candidate identified). The Melanoma sample is annotated (taken from original publication) with black as melanoma, red as stroma and yellow as immune infiltrate. A cutoff of 0.0 was used when visualizing the TLS-score predictions.

**Supplementary Table 2.** Genes included in TLS-signature.

| Gene | Coefficient | Gene | Coefficient | Gene | Coefficient |
| --- | --- | --- | --- | --- | --- |
| MS4A1 | 0.000595692124909 | CD69 | 0.000178523456895 | ALG8 | 0.000144285394958 |
| B2M | 0.000332672042219 | SYT11 | 0.00017618286567 | CD79A | 0.000143369081063 |
| TRBC2 | 0.000317378385003 | FYB | 0.000175798252131 | UHRF1BP1L | 0.000143360448522 |
| LTB | 0.000314349966323 | CD48 | 0.000175141314241 | MDFIC | 0.00014292847454 |
| TIAM1 | 0.000314177344547 | PLA2G2D | 0.00017383613927 | TARBP1 | 0.000142217292012 |
| ZC3H12D | 0.000313029236991 | SPIB | 0.00017356648257 | MICB | 0.000142213802666 |
| SCML4 | 0.00030351675486 | LIMD2 | 0.000172949993333 | B3GLCT | 0.000142087554698 |
| CXCR4 | 0.000295575363742 | ARHGAP15 | 0.000171935625146 | CASP2 | 0.000141827504695 |
| CD19 | 0.000288765012981 | BCL11B | 0.00017006201514 | SELL | 0.000141554153463 |
| TRAF3IP3 | 0.000285710237725 | AKNA | 0.000169636422625 | EMB | 0.000141385335943 |
| BTG1 | 0.000271035711673 | TNFAIP8 | 0.000168196592318 | CD37 | 0.000141033972406 |
| BIRC3 | 0.000270754219697 | CD2 | 0.000167327817843 | ABCD4 | 0.000140912625775 |
| CD52 | 0.000263835835808 | DDX5 | 0.000166677867478 | AP4B1 | 0.000140867027941 |
| RAC2 | 0.000257684640648 | POGLUT1 | 0.000166524246914 | TTBK2 | 0.000140579746916 |
| CD83 | 0.000251490238498 | TIPARP | 0.000165554981735 | FAM65B | 0.000140571174096 |
| HLA-B | 0.000249477062823 | GAREM2 | 0.000165279736338 | TMEM5 | 0.000140404709944 |
| PTPRC | 0.00024628671493 | EPHA4 | 0.000165144835794 | ANKDD1A | 0.000140393391408 |
| TRAC | 0.000243828273295 | IL2RB | 0.00016475163977 | WIPF1 | 0.000140275511107 |
| SLA | 0.000239561152957 | GRPEL2 | 0.000164137100584 | NKRF | 0.000140206314254 |
| CXCL9 | 0.00023484222349 | ADAM28 | 0.000163998734444 | SEPT6 | 0.000140109149509 |
| CORO1A | 0.000231949711471 | VWA8 | 0.000163958977584 | IL10RA | 0.000139939044216 |
| IL16 | 0.000227806544677 | FAR1 | 0.000163316470036 | SLC25A33 | 0.000139753022653 |
| CYTH1 | 0.000222133222142 | TMSB4X | 0.000162675011343 | LMO2 | 0.000139663150449 |
| ZFP36L2 | 0.00022125659097 | CACNA1I | 0.000161818137099 | DUSP2 | 0.000139263146875 |
| ITGAL | 0.000220774848163 | SMCHD1 | 0.000160065486459 | NACA | 0.000138814706626 |
| ICOS | 0.000220531627722 | HLA-DQB1 | 0.000159989878407 | SCIMP | 0.000138632593831 |
| FAM96B | 0.000219830644763 | LNPEP | 0.000158710432572 | ZSCAN29 | 0.000138317307187 |
| CXCL13 | 0.000219689264426 | RCHY1 | 0.000158638904959 | RNASET2 | 0.000138178147064 |
| PASK | 0.000216719757693 | GPSM3 | 0.000157818119632 | RFFL | 0.000138106631077 |
| CCR7 | 0.000211367390464 | TRBC1 | 0.000157724735433 | CD96 | 0.000138037067735 |
| SNX13 | 0.000210712369247 | TSC22D3 | 0.000156868044902 | CD79B | 0.000135989992381 |
| TRAF5 | 0.000209622341485 | DOCK8 | 0.000156646041763 | DGKE | 0.000135961951047 |
| LRMP | 0.000208922706969 | SPOCK2 | 0.000156272915968 | SGSM2 | 0.000135605121307 |
| BLK | 0.000208263281006 | PIP4K2A | 0.00015617577339 | FN3KRP | 0.000135460976542 |
| IL7R | 0.000207819468811 | CCDC50 | 0.00015532079662 | ZNF700 | 0.000135343521919 |
| PRKCB | 0.000206018165061 | GZMK | 0.000154890466818 | TBC1D23 | 0.000135309288028 |
| IL11RA | 0.00020215370056 | CPSF7 | 0.00015484366527 | PIK3IP1 | 0.000135100371679 |
| IL2RG | 0.000200475332668 | CD53 | 0.000154433698911 | SP140 | 0.000134814720212 |
| CD247 | 0.000200350709432 | SLAMF6 | 0.000153225703526 | DEF6 | 0.000134762514514 |
| QRICH1 | 0.000199184724442 | BARX2 | 0.000152592481906 | EVI2A | 0.00013474131392 |
| CARMIL2 | 0.00019838434595 | FUCA1 | 0.000151902661181 | CARD11 | 0.000134728651734 |
| C5orf56 | 0.000196288725541 | CCL19 | 0.000151759813087 | HNRNPA1 | 0.000134434803067 |
| CD3E | 0.000196279783133 | ACAP1 | 0.000151533777593 | RNF43 | 0.000133758771941 |
| DNMBP | 0.00019550043764 | IBTK | 0.000151216643474 | HLA-DMB | 0.000133751800931 |
| RHOF | 0.00019001882019 | OSBPL7 | 0.000151164146772 | CD5 | 0.000133081864036 |
| ARHGEF1 | 0.000189847012439 | STK17A | 0.000149839180989 | TBC1D10C | 0.000132825037694 |
| KLF12 | 0.000188405291697 | CD3G | 0.000149756396866 | SEPT1 | 0.000132610015045 |
| TXNIP | 0.000187362619779 | ATP8A1 | 0.000149099799003 | ZBTB46 | 0.000132412254372 |
| LYZ | 0.000187060745086 | KLF2 | 0.000149027298158 | S100PBP | 0.000132239263083 |
| TCL1A | 0.0001870026952 | CD27 | 0.000148457414886 | EIF4H | 0.000132006812558 |
| SAFB | 0.000186077044672 | COA1 | 0.000148200697177 | NELL2 | 0.000131969279709 |
| DGKA | 0.000185571574722 | CXCR5 | 0.000148114866973 | TMED8 | 0.000131946332862 |
| BRE | 0.000183349162999 | EPB41 | 0.000147422991384 | MLLT10 | 0.000131921796035 |
| PLEKHG1 | 0.000183107764966 | GTF2E1 | 0.000146664134548 | CASP1 | 0.000131851115526 |
| PTPN22 | 0.000182056748811 | SLAMF1 | 0.000146196045923 | TESPA1 | 0.000131262931785 |
| LAMP5 | 0.000181972075947 | TMC6 | 0.000145610417408 |  |  |
| ZNF558 | 0.000181232416102 | UBA7 | 0.000144881439034 |  |  |
| JAK3 | 0.000179071216954 | MSN | 0.000144873617036 |  |  |

### File Listing (Supplementary Data)

- **Supplementary Data 1** CORESIG.xlsx - genes included in each of the respective core signatures (2 immune associated and 1 tumor associated)
- **Supplementary Data 2** CLUSTERANNOT.xlsx - cluster annotations and the motivations behind these
- **Supplementary Data 3** MARKERGENES.xlsx - marker genes for each of the expression based clusters (within each patient)
- **Supplementary Data 4** RINGGENES.tsv - list of genes that were excluded in parts of the analysis due to their aberrant behavior; forming ring-like patterns in several capture areas.
- **Supplementary Data 5** CORRS.zip - correlation matrices for all three tiers. Matrices are sorted in subfolders according to their tier, naming follows the convention: corrs-{tier}/{tier}-{samples}-corr-plot.png
- **Supplementary Data 6** PATHWAYS.zip - enriched pathways for each of the expression based clusters. Naming follows the conventions : {patient}_GO_BP.xlsx
- **Supplementary Data 7** STSCPROP.zip - Proportion estimates across all tiers and samples. Naming follows the convention : {tier}/{sample}-proportion.tsv
- **Supplementary Data 8** STSCVISUAL.zip - Visualization of proportion estimates across all tiers and samples. Naming follows the convention : {tier}/{sample}-proportion-visual.png.
- **Supplementary Data 9** STSCENR.zip - enrichment/depletion results for all cell types across all tiers. Naming follows the convention {annotation}/{tier}/{sample}-enrichment.png, with annotation = [cluster,pathologist]x.
- **Supplementary Data 10** CLUSTERENR.zip - enrichment of expression-based clusters and the regions defined by the pathologist. Naming follows the conventions {sample}_cluster_vs_annotation_overlap.xlsx
- **Supplementary Data 11** STSCGENELIST.tsv - genes included from the single cell data when running *stereoscope*.
- **Supplementary Data 12** TLSENR.tsv - enrichmed pathways when subjecting the TLS-signature to functional enrichment analysis (GO:BP)
- **Supplementary Data 13** TLSCOEF.tsv - coefficient for all genes obtained upon fitting the linear model (trying to predict TLS-score) to patient G and H.

